# Pathogenic O-GlcNAc dyshomeostasis is associated with cortical malformations and hyperactivity

**DOI:** 10.1101/2025.04.27.650822

**Authors:** Florence Authier, Asad Jan, Islam Faress, Christian Stald Skoven, Iria Esperon-Abril, Shagana Tharmakulasingam Balasubramaniam, Kévin-Sébastien Coquelin, Jens R. Nyengaard, Carsten Scavenius, Benedetta Attianese, Oscar G. Sevillano-Quispe, Simon Fristed Eskildsen, Jesper Skovhus Thomsen, Brian Hansen, Daan M. F. van Aalten

## Abstract

Missense variants in the O-GlcNAc transferase (*OGT)* gene have recently been shown to segregate with a syndromic form of intellectual disability (OGT-ID), underscoring the importance of protein O-GlcNAcylation in brain function. However, the underlying pathophysiological mechanisms linking ID to potential OGT malfunction—whether developmental, neurophysiological, or both—remain unclear. Here, we present comprehensive analyses encompassing behaviour and brain architecture of a rodent model carrying the pathogenic C921Y OGT-ID variant. These mice show a range of behavioural deficits, including hyperactivity, impulsivity, and associative learning phenotypes. Structural studies, using micro-computed tomography and magnetic resonance imaging, revealed reduced skull size, microcephaly, reduced cortical thickness and hypoplastic corpus callosum. Detailed histological analyses revealed dysplastic changes in the neocortex, predominantly affecting the superficial layers of cingulate cortex. Mechanistically, quantitative proteomic analyses revealed O-GlcNAc dyshomeostasis associated with distinct perturbed molecular pathways involved in brain development. Taken together, these data reveal neurodevelopmental defects associated with O-GlcNAc dyshomeostasis and provide a platform for dissecting mechanism and treatments of OGT-ID.

## Introduction

Intellectual disability (ID) is a prevalent neurodevelopmental condition estimated to affect 1-3% of the global population and is characterized by a significant impairment in cognitive function, along with compromised adaptive and social behaviour^1^. Developmental anomalies in the human cerebral cortex, particularly the neocortex, are common causes of intellectual disability (ID)^2,3^. A recently reported syndromic form of ID is associated with missense variants in the *OGT* gene, which resides on chromosome Xq13.1, and encodes the O-linked N-acetylglucosamine transferase (OGT). This enzyme facilitates the covalent attachment of O-linked b-N-acetylglucosamine (*O*-GlcNAc) to the hydroxyl groups of serine and threonine (Ser/Thr) residues on nuclear, cytoplasmic and mitochondrial proteins^4,5^. *O*-GlcNAcylation is a dynamic and highly conserved modification governed solely by two enzymes, OGT for attachment^6,7^ and O-GlcNAcase (OGA) for removal^8^. This sets O-GlcNAcylation apart from other post-translational modifications, such as phosphorylation and ubiquitination, that rely on hundreds of “writer” and “eraser” enzymes, associated with specific sets of “reader” proteins^9,10^. O-GlcNAcylation regulates cellular functions, such as transcriptional activation^11^, gene expression^12^, stress response^13^ and proteostasis^11^, in response to physiological changes. Structural studies of OGT have demonstrated that the enzyme is composed of an N-terminal tetratricopeptide repeat (TPR) domain, involved in substrate recognition and binding of OGT interactors^9,10^, and a catalytic glycosyltransferase domain. In addition to catalysing O-GlcNAcylation, the catalytic domain is involved in the proteolytic activation of the transcriptional coregulator Host Cell Factor 1 (HCF-1), itself an ID-associated protein^14,15^.

OGT and OGA are notably abundant in the brain, with particularly enrichment in hippocampal formation and related structures^16^. Thus, O-GlcNAcylation plays a critical role in neuronal survival, development, and synaptic function. For instance, pan-neuronal knockout of OGT results in severe neurodevelopmental defects and early neurodegeneration^17,18^. Moreover, deletion of OGT in the dopaminergic neurons of substantia nigra or peripheral neurons in the dorsal root ganglion induces widespread apoptosis^19,20^. At the synapse, OGT and OGA display distinct localization patterns: OGA is absent from the post-synaptic density, while OGT is distributed evenly between pre- and post-synaptic compartments in excitatory synapses^21,22^. In this context, aynaptosome mass spectrometry (MS) data have revealed that nearly 20% of synaptic proteins, including ankyrin G, CaMKIV, and GluA2, are modified by O-GlcNAc, which dynamically regulates excitability and transmission^23,24^.

To date, 17 pathogenic missense variants in the *OGT* gene have been identified as causal for a newly described syndromic form of ID, termed OGT-ID, also known as O-GlcNAc Transferase Congenital Disorder of Glycosylation (OGT-CDG)^25–29^. This syndrome is frequently comorbid with neurological and psychiatric disorders such as epilepsy and autism spectrum disorder (ASD), both of which are associated with focal cortical dysplasia (FCD)^30–32^. In addition to ID and maladaptive behaviour, OGT-ID patients exhibit a plethora of muscular, facial and neurological abnormalities, such as hypotonia, craniofacial dysmorphia, microcephaly, fifth finger clinodactyly, and developmental delay^25,26,33^. Missense and exon-skipping variants in OGT have been found across both functional domains (i.e. TPR and catalytic), with affected patients exhibiting similar clinical features. These observations hint towards pathogenic mechanisms whereby the OGT variants affect neurodevelopmental processes and/or brain function leading to intellectual disability. Nevertheless, understanding the potential impact of OGT-ID variants on brain development has remained challenging, primarily due to the lack of viable vertebrate models since germline knockout (KO) of *Ogt* in mice leads to embryonic lethality^34^.

Among OGT-ID variants, the catalytically impaired C921Y variant has been the most extensively studied. It was identified in three affected brothers born to a healthy, non-consanguineous couple in Denmark. The affected individuals exhibit developmental delay, autistic features, dysmorphic traits, osteoporosis and seizures^35^. Functional studies have shown that the OGT^C921Y^ variant has decreased glycosyltransferase activity, both *in vitro* and in mouse embryonic stem cells (mESCs). In OGT^C921Y^ mESCs, the mutation reduces O-GlcNAcylation and decreases expression of stem cell markers Oct4, Sox2, and alkaline phosphatase (ALP)^32^, suggesting that OGT plays a critical role in embryonic stem cell self-renewal and pluripotency^32^. In *Drosophila melanogaster* models, the OGT^C921Y^ mutation has been shown to reduce O-GlcNAcylation during development^36^, a defect that can be rescued by genetic or pharmacological inhibition of OGA. This mutation disrupts larval neuromuscular junction development and shortens sleep bout duration, with both effects being partially reversible through OGA inhibition. These findings suggest that certain aspects of OGT-ID pathology are developmental in origin, while others may be reversible. Although these studies indicate that early differentiation and development are affected, a vertebrate model is required to dissect the developmental and brain-wide effects of OGT-ID variants, as the precise mechanisms remain unknown.

Here, we use an OGT^C921Y^ mouse model of OGT-ID to reveal a range of behavioural deficits, including hyperactivity, impulsivity, and associative learning phenotypes. Structural studies, using micro-computed tomography and magnetic resonance imaging, revealed reduced skull size, microcephaly, reduced cortical thickness and hypoplastic corpus callosum. Detailed histological analyses uncovered nodular cortical dysplasia, predominantly affecting the superficial layers of cingulate cortex. Mechanistically, quantitative proteomic analyses suggested perturbed regulation of distinct perturbed molecular pathways involved in brain development in the presence of O-GlcNAc dyshomeostasis. Taken together, these data reveal neurodevelopmental defects associated with O-GlcNAc dyshomeostasis and provide a platform for dissecting mechanism and treatments of OGT-ID.

## Materials & Methods

### Animal husbandry

The OGT^C921Y^ line was previously generated and reported^37^ and was maintained under C57BL/6J background (Janvier, France). Animal cohorts were obtained from crossing of male OGT^WT^ with female OGT^C921Y/+^. Only male mice were used in all experiments. Animals were housed in digitally ventilated cages (Tecniplast, Italy) with water and food available *ad libitum* and 12/12 h light/dark cycles in the Skou animal facility of Aarhus University. All animal studies and breeding were performed in accordance with the ARRIVE guidelines and the European Communities Council Directive (2010/EU) and were approved by the Danish Animal Experiments Inspectorate (Dyreforsøgstilsynet), under Breeding license 2022-15-0202-00135 and Project licenses: 2023-15-0201-01426 and 2020-15-0201-00421.

### Non-invasive monitoring of spontaneous activity in home cages (DVC)

Patterns of locomotion and spontaneous activity were continuosly monitored over 24 hours in home cages through a specialized digitally ventilated cages (DVC) platform (Tecniplast, Italy) between the age of 30 days and 105 days (termination). This platform is based on electrical capacitance sensing technology, with incorporates a sensor board equipped with an integrated circuit comprising 12 electrodes directly beneath the floor of the cages^38^. The DVC circuit measures changes in the electrical capacitance signal from each electrode in response to the movement of a water-filled body (animal) close to or away from a given electrode. The measurements, performed approximately 4 times per second, are remotely relayed to the centralized DVC analytics platform (Tecniplast, Italy). In this web-based interface, time-stamped data for each cage can be visualized using in-built tools (e.g. daily rhythms, cumulative activity/locomotion index aggregated per minute/hour/day, bedding status, light or dark period activity, heatmaps etc.). In the default setup, the DVC analytics web-interface plots the animal locomotion index as arbitrary units normalized between 0% and 100%, representing the overall activity performed in the cage by the animals, i.e., the signal is measured for each cage and not each animal.

### Behaviour assessment

Prior to behavioural testing, all animals were handled daily by the experimenter for one week. All handling took place in the experimental room with an ambient light setting of 25-30 Lux. All animals were housed in groups of 2 to 3 individuals. Animals were 10 weeks old at the start of behavioural testing, reaching 14 weeks old at the end. Before any behavioural test, mice were place in the experiment room for at least 30 min prior to testing. The data analyses and video quantification were performed blindly with respect to the genotype. The sample size was determined based on previous experience and validated using Post-hoc Power Calculation using ClinCalc online tool (https://clincalc.com/stats/Power.aspx).

### Locomotor behaviour and sensorimotor coordination

*Open Field:* The activity of mice in an open field maze was recorded using ANY-maze video tracking software (Stoelting Europe, Ireland). Individual animals were placed in a 33.5 x 33.5 x 39 cm opaque box and were allowed to explore for 10 min over three consecutive days. Time spent in the centre and periphery and time moving in the periphery were analysed to investigate parameters of locomotion and anxiety.

*Rotarod:* Coordination skills were assessed using rotarod apparatus. Mice were placed on a Ugo Basile NB 80534 rotarod with an increase in speed from 4 to 40 rpm and an acceleration time of 2 min (40 rpm/min is reach after 2min) for maximum 5 min. Latency to fall from the rotarod and speed were measured.

*Static rods*: Mice were placed at the extremity of the rod with head facing the void. Time spent by the mouse to perform a t-turn and to reach the goal platform were recorded. If the animal fall <5 s (presumably due to a misplacement), the mouse was tested again. After three consecutive fails or an up-side down of a mouse, a 120 s score was reported. Five rod diameters (35, 28, 22, 15 and 10 mm) were used for each animal.

*Pole test*: Mice were placed close to the top of a pole (40 cm) with the head of the mouse facing up. Times to perform a t-turn and reach the ground were recorded. The test was performed three times for each mouse. If the mouse succeeded the three attempts, a score of 3 was reported. For each slide and fail to perform the test within the 120 s cut off, an additional 1 point was incremented in the total score.

### Cognitive performance

*Novel Object Recognition*: Novelty associated short- and long-term memory were assessed in an open-field maze. During the familiarisation phase mice are free to explore the maze containing two identical familiar objects for 10 min. After 90 min for short-term and 24 h for long-term memory, one of the familiar objects is replaced by a novel object and mice are free to explore the maze for 10 min. Time spent exploring objects and number of explorations was measured until reaching a total exploration time of 20 s for each mouse. *Spontaneous alternation:* The spontaneous alternation test was performed in a T-maze and was used to assess spatial working memory. Mice were free to choose left or right arms for 7 trials. Each arm entry was recorded to calculate the percentage of spontaneous alternation corresponding to the number of correct Left-Right (L/R) or Right-Left (R/L) sequences. Mice that were able to remember which arms they had entered most recently would choose a different one to explore.

*Aversive conditioning and recall*: The apparatus consisted of an open-top cage (24 × 20 × 30 cm) with metal floor bars, placed inside a soundproof cubicle (55 × 60 × 57 cm) (Ugo Basile, Italy). Three minutes after being placed in the conditioning chamber, the mice were conditioned using four tone-foot shock (CS-US) pairings (n = 9 per group). Each pairing consisted of a 25 s, 7 kHz tone (CS), followed by a 25 s gap (trace period), and then a 1.5 s foot shock (US) at 0.5 mA. After the conditioning session, the animals were isolated for 10–15 min before being returned to their home cage with their littermates. Long-term memory recall was assessed in a novel context 24 h after conditioning. Following a 3 min acclimatisation period in the novel context, the mice were exposed to four CS presentations without foot shocks. The intertrial interval ranged from 120 to 180 s for both the conditioning and testing sessions. The behavioural responses were recorded using a top-mounted camera and freezing and locomotor activity were automatically scored using ANY-maze software (Stoelting Europe, Ireland). The freezing percentage represents the time the mouse spent freezing during the CS presentation or the total duration of the time bin.

### Anxiety behaviour

*Elevated-plus maze:* Anxiety-like behaviour was assessed using the elevated plus maze paradigm. Mice were placed in the centre of the cross-shaped maze comprised of two open arms (125 Lux) and two closed arms (25 Lux) and allowed to explore the maze for 5 min. Time, distance and number of entries in each of the open and closed arms were analysed as an approximation for anxiety.

*Dark-light paradigm*: Light-like anxiety behaviour was assessed using the dark-light paradigm at 180 Lux. Mice were placed in an arena containing both dark and light compartments. At the start of the test, mice were placed in the dark compartment and were free to explore both compartments for 10 min. Time, distance and number of entries in the light compartment were analysed to investigate the level of anxiety.

### Compulsive behaviour

*Marbles*: Before the test, 20 glass marbles (1.6 cm in diameter) were placed on the top of the bedding (in five rows of four marbles) in a cage with 5 cm bedding. At the end of the test, the number of buried marbles (covered by at least 75% of bedding) was counted.

*Digging and self-grooming:* Mice were placed in a cage with 5 cm (digging) or 1 cm (self-grooming) bedding for 3 min. Number of digging/self-grooming/rearing events and time spent digging/self-grooming/rearing were recorded. *Nesting:* Mice were placed in a cage with 1 cm bedding in the presence of a cotton pad as nesting material for 1 h. Cotton pads were weighed before and 24 h after testing to allow drying. Percentage of material removed was quantified.

*Litter burrowing:* Mice were placed in a cage in the presence of a PVC tube filled with 120 g of litter bedding for 30 min. Percentage of litter material removed was quantified.

### Brain perfusion for structural analyses

Brains were perfusion fixed and prepared for in-skull MRI (OGT^WT^ n = 15 and OGT^C921Y^ n = 16). Perfusion fixation was performed after the mice had been anesthetized by an intraperitoneal injection of Euthanimal (250 mg/kg, Alfasan, 088672). Then, the brain was fixed by transcardiac perfusion at 125 mmHg, to be close to physiologic brain perfusion as previously reported^39^ using 25 mL heparinized (Heparin, 0.2 mL/100 mL, 5000 IU/mL, Pan Pharma, 482480) Natriumchlorid (9 mg/mL, B Braun, 5/389885/0417) for 3 min 30 s, followed by 25 mL of buffered 10% formalin solution (VWR Qpath Chemicals, 11699404) for 3 min 30 s. After decapitation, the mandible and extracranial tissue were removed from the skull to avoid susceptibility artifacts from air bubbles trapped in fur and cavities during imaging. Hereafter, the in-skull brains were stored in 10% formalin solution for at least one week prior imaging. One WT mouse was excluded due to misperfusion.

### Magnetic resonance imaging (MRI) and image analysis

Before imaging, the fixed brain samples were washed in PBS for at least 24 h to increase MRI signal by removal of excess fixative^40^. For imaging, the samples were subsequently mounted in a 15 mL centrifuge tube filled with a perfluorocarbon based liquid (Fluorinert, 3M, PN: FC-770) as is standard^41–46^.

MRI data collection: MRI scans were acquired on a 9.4 T preclinical system (BioSpec 94/20, Bruker Biospin, Ettlingen, Germany) using a bore-mounted 25 mm quadrature transmit-receive coil. To avoid sample vibrations, the tube containing the sample was secured in a custom polyethylene foam cylinder inside the coil. In-house 3D-printed sample holders ensured consistent positioning of the samples throughout experiments. High-resolution B0 maps were acquired before each sequence, allowing shimming using Bruker’s MAPSHIM. Both DKI data and structural data were acquired for each sample.

Diffusion kurtosis analysis: DKI data was collected using an 8-segmented diffusion-weighted spin-echo EPI sequence with a 150 × 150 μm in-plane resolution and 250 μm slice thickness (60 slices for whole-brain coverage). Five unweighted volumes were acquired for signal normalization followed by 30 isotropically distributed encoding directions at each of three non-zero b-values (0.5, 1.0, 2.0 ms/μm^2^). Additional scan parameters were time between diffusion gradients (Δ) = 15 ms, diffusion gradients duration (δ) = 6 ms, 20 averages, effective echo time (TE) = 27.7 ms, repetition time (TR) = 3500 ms, bandwidth = 278 kHz, resulting in a DKI scan time of 19h26m40s per animal. In addition, a rapid acquisition with relaxation enhancement (RARE) sequence with a 50 × 50 μm in-plane resolution and 250 μm slice thickness was performed. Here, the 60 slices were positioned identically to the DKI data to allow for precise multi-atlas segmentation (MAS, details below) and ROI-specific extraction of DKI parameters. The scan parameters used were effective TE = 10.5 ms, TR = 3000 ms, 30 averages, and RARE factor = 2, with a scan time of 2h28m30s per animal. For the volumetric analysis, we acquired data with an isotropic resolution of 50 μm using a 3D fast low-angle shot (FLASH) sequence. Scan parameters for this were: TR = 88.5 ms, TE = 13.7 ms, matrix size = 360 × 198 × 300, FOV = 18 × 9.9 × 15 mm, and 4 averages, resulting in a scan time of 6h31m21s per animal. For all scan types, data quality was ensured by visual inspection and samples rescanned if needed to ensure consistently high data quality for subsequent analyses. For diffusion kurtosis (DKI) analysis all DKI data from all samples were pre-processed in MATLAB (MathWorks Inc., v. 2022a) for noise floor correction, denoising and Gibbs ringing removal as described in^46,47^. After pre-processing, DKI data analysis was performed using inhouse MATLAB scripts as previously described^47,48^ yielding metrics of mean water diffusivity (MD), tissue anisotropy (FA) and the mean kurtosis (MK, an index of tissue microstructure) in each voxel^47^.

*Multi-atlas segmentation*: To systematically extract regional information from the mouse brain, a multi-atlas segmentation (MAS)^46,49,50^ was performed using 10 ex vivo NeAt templates from C57/BL6J mice (60,61). This followed procedures as described previsouly^46^. Thus, high-resolution labelled images were obtained and then down sampled to match the in-plane resolution of the DKI data allowing extraction of DKI metrics from anatomically well-defined regions of interest (ROIs). Regional voxel values of DKI metrics were filtered for outliers (defined as values exceeding 3 times the median absolute deviation) and used for statistical analysis.

*Volumetric analysis*: The high-resolution FLASH images were processed using an in-house pipeline^46^ applying B1 inhomogeneity correction^53^, denoising^54^, and intensity normalization. Spatial alignment with a high-resolution template of the C57BL/6J mouse^55^ was done by manually initializing a linear registration^56^ followed by a non-linear registration^57^. Neuroanatomical labels from the C57BL/6J mouse atlas were subsequently transformed and resampled to data native space using the calculated deformation fields and affine transformations for calculation of individual regional volumes. From this, absolute brain volume and regional volumes were calculated. For all brains, relative regional volume (RRV; region size as % of total brain volume) was calculated to account for total brain size variation.

*Statistics*: To investigate group differences, permutation tests were performed for either total brain volume (1M permutations), regional volumes (40 regions, 100k permutations) or extracted voxelwise DKI metrics for each region (20 regions, 100k permutations). Uncorrected significance is reported for *p* values < 0.05 and indicated by asterisks (*). Pound symbols (#) indicate significance below α = 0.05 divided by total number regions tested for group mean (green) and group median (red), respectively. Section signs (§) indicate significance (*p* < 0.05) with *p* values adjusted for false discovery rate (Benjamini-Hochsberg)^58^.

*Cortical thickness*: Cortical thickness was calculated as previously described^59^. Briefly, Laplace’s equation, with fixed boundary conditions for each of the inner and outer surfaces, was solved. For this, the inner and outer surfaces of the cortex were defined based on the anatomical atlas and transformed to the given mouse. For each point on the cortical surface, the length of a streamline connecting the inside and outside surfaces was used to define the thickness. Cortical thickness was averaged within the bilateral frontal, occipital, and parieto-temporal lobes as well as the entorhinal cortex. Statistical maps of group differences in cortical thickness were generated by fitting a general linear model at each surface vertex (SurfStat, http://www.math.mcgill.ca/keith/surfstat/). Given the multiple comparisons performed, statistical maps were family-wise error (FWE) corrected using random field theory [38] with α = 0.001 as cluster defining threshold. All statistical maps were thresholded at *p* = 0.05 (uncorrected and corrected).

### Micro Computated Tomography (MicroCT)

Following MRI, the in-skulls brain samples were imaged using MicroCT scanning (vivaCT 80, Scanco Medical AG, Brüttisellen Switzerland). Skulls of 20 week old mice were placed in the scanner and imaged using 500 projections over 180°, an isotropic voxel size of 39 µm, X-ray voltage of 55 kVp, X-ray current of

105 µA, and an average time of 200 ms. Images were reconstructed and converted to DICOM files that were exported for subsequent analysis in 3D slicer (http://www.slicer.org) and rendered in 3D. Skulls were isolated applying a threshold that separates bone from soft tissue. Segmented skulls and extracted endocasts were exported as 3D models. Furthermore, the coordinates of 45 surface landmarks were registered in a semi-automated fashion: a random skull was chosen as template model and landmarked manually, and these landmarks were automatically applied to the rest of the dataset using the ALPACA module in 3D Slicer. Additionally, the PseudoLMGenerator module was used to obtain a dense network of surface landmarks consisting of 768 points from the skulls and used as input for Principal Components Analysis (PCA). The image processing was performed with the operator blinded for the group distribution. For 3D visualization of differences between WT and OGT^C921Y^ mice by heat maps, models for the average shapes of OGT^WT^ and OGT^C921Y^ skulls were generated from the General Procrustes Analysis (GPA) module, aligned, superimposed, and the model-to-model distance was measured. Skull shape and size differences were assessed by measuring distances between the 45 surface landmarks and performing Euclidean distances matrix analysis. A previously collected cohort of 8 weeks old mice skulls were also analysed with this pipeline^60^. One WT skull was excluded from analysis due to imaging artefact.

### Histology and immunofluorescence (IF) microscopy

Following MicroCT scanning, formalin fixed and paraffin embedded brain sections (10 µm thickness) were obtained from the male WT and OGT^C921Y^ mice (n = 6 per group). To better characterize the radial and tangential cytoarchitecture in the neocortex, as well as to assess the bilaterality of any incidental findings in brain structure, one hemisphere was cut in sagittal orientation while the opposite hemisphere was cut in coronal orientation. Sections were deparaffinised and stained with haematoxylin and eosin, as described^61^. Serial sections were stained with cresyl violet for assessing the cellular arrangement of neurons in cortical layers, and separately with Luxol Fast blue for visualizing white matter distribution, including the arrangement of large tracts^61^.Then, high resolution views were obtained with an Olympus VS120 digital slide scanner equipped for bright field imaging. Slide scans were imported into Qupath (v. 0.5.1)^62^ and regions of interest (ROI) were outlined manually guided by the Mouse Brain Atlas (Paxinos and Franklin’s The Mouse Brain in Stereotaxic Coordinates, 4th Edition). ROI were segmented using the Qupath cell detection on the haematoxylin channel.

Cellular identity was further verified by immnuofluorescence (IF) microscopy using the following primary antibodies: anti-NeuN (1:500; ABN78; Sigma-Aldrich) and anti-glial fibrillarly acidic protein- GFAP (1:500; ab68428; abcam). For this purpose, sections were deparaffinised and incubated in a blocking buffer comprising 5% normal goat serum in Tris-buffered saline for 1 h at room temperature. Then, the sections were incubated overnight at 4°C with the primary antibodies diluted in phosphate-buffered saline containing 0.3% Triton-X and 0.5% bovine serum albumin- BSA: IF detection was performed by fluorophore conjugated secondary antibodies: Alexa-Fluor488 goat anti-rabbit (1:1000; A-11034; Thermo Fisher) and Alexa-Fluor594 goat anti-mouse (1:1000; A-11005; Thermo Fisher). Image acquisition was performed on a Leica DM6000 upright microscope equipped with a fluorescence light source. Multiplex IHC analyses were performed in Discovery Ultra automated staining system (Roche Diagnostics) after deparaffinization and heat-induced antigen retrieval using DISCOVERY CC1 buffer (Roche Diagnostics catalogue # 06414575001) for 32 minutes at 95 °C. Then, sections were sequentially incubated with the following primary antibodies: anti-NeuN (Rabbit polyclonal, Sigma-Aldrich catalogue # ABN78; dilution, 1:500), anti-Olig2 (clone: EP112; Rabbit monoclonal, Roche Diagnostics catalogue # 07667973001; dilution, 1:500), anti-Reelin (Goat polyclonal, Novus Biologicals catalogue # AF3820, dilution: 1:100) and anti- Pou3F2/BRN2 (Rabbit polyclonal, Proteintech catalogue # 14596-1-AP; dilution, 1:250). Chromogenic detection was performed using Discovery Omnimap Horseradish peroxidase (HRP)-conjugated secondary antibodies with following chromogens: DISCOVERY Teal HRP (for NeuN; Roche Diagnostics catalogue # 08254338001), DISCOVERY ChromoMap DAB (for Olig2: Roche Diagnostics catalogue # 05266645001), DISCOVERY Purple (for Reelin; Roche Diagnostics catalogue # 07053983001) and DISCOVERY yellow HRP (for POU3F2; Roche Diagnostics catalogue # 8502641001). In order to avoid cross-reaction between antibodies, denaturation between chromogenic detection of markers was performed using ULTRA CC2 buffer (Roche Diagnostics catalogue # 05424542001) at 100 °C for 24 minutes. High resolution views were obtained with an Olympus VS120 digital slide scanner in bright field imaging mode.

### Tissue collection and dissociation for biochemical analyses

The prefrontal cortex was rapidly isolated from whole 16 week old male mouse brain (n = 3 per group), snap frozen and stored at −80 °C until processing. Tissues were disrupted in phosphate-buffered saline (PBS) two times at 5000 rpm for 30 s with 10 s break using a Precellys^®^ 24 Touch homogenizer (Bertin Technologies). Homogenates were split in half for further protein and RNA extractions.

### Mass Spectrometry (MS) and data analysis

Protein extracts from prefrontal cortex tissues were prepared for mass spectrometry using S-Trap micro spin columns (Protifi), including three washes with 50% CHCl₃/50% MeOH. Trypsin digestion (Proteomics grade, Sigma-Aldrich) was performed for 16 h at 37 °C. The resulting peptides were lyophilized and dissolved in 0.5% formic acid. LC-MS/MS was conducted using an EASY-nLC 1200 system (Thermo Scientific) connected to an Orbitrap Eclipse Tribrid Mass Spectrometer (Thermo Scientific) with a 2 cm trap column (100 μm i.d.) and a 15 cm analytical column (75 μm i.d.), both packed in-house with ReproSil-Pur C18-AQ 1.9 μm resin (Dr. Maisch GmbH). Peptides were eluted at 250 nl/min using an 80 min gradient from 5% to 44% phase B (0.1% formic acid and 80% acetonitrile), followed by a 30 s gradient to 100% phase B and 5 min at 100% B. Protein identification and quantification were performed using Proteome Discoverer 2.5 (Thermo Scientific). Data were searched against the mouse reference proteome (uniprot.org) using the Sequest search engine with the following parameters: MS error tolerance of 10 ppm, MS/MS error tolerance of 0.02 Da, trypsin as the protease with two missed cleavages, and carbamidomethylation as a fixed modification. Variable modifications included HexNAc (ST) and oxidation (M). Label-free quantification was based on precursor ions using unique peptides quantified in at least 2 out of 3 replicates. Peptide intensities were normalized to total peptide intensity and scaled using the average of all samples. Protein ratios were based on summed peptide abundances with imputation using replicate-based resampling. Significantly regulated proteins were identified using ANOVA, with adjustments for multiple testing.

### Western immunoblotting

Brain homogenates were lysed using 10x RIPA buffer (Cell Signaling) as previously described^37^. For MS, 50 mL of lysates were stored at −80 °C until further processing. For western blot, the rest of the lysates were centrifuged at 14,000 rpm for 20 min at 4 °C, and the protein concentration was determined with Pierce^TM^ BCA Protein Assay kit (Thermo Scientific, 23227). Proteins (20 μg) were separated on precast 4-12% NuPAGE Bis–Tris Acrylamide gels (Invitrogen) and transferred to nitrocellulose membrane. Membranes were incubated with primary antibodies in 5% bovine serum albumin in Tris-buffered saline buffer with 0.1% Tween-20 overnight at 4 °C. Anti-OGA (1:1000 dilution; HPA036141; Sigma), anti-O-GlcNAc (RL2) (1:1000 dilution; NB300-524, Novus Biologicals), anti-OGT (F-12) (1:1000 dilution; sc-74546; Santa Cruz), mouse anti-actin (1:5000 dilution; A5441; Merck) antibodies were used. Next, the membranes were incubated with IR680/800-labeled secondary antibodies at room temperature for 1 h. Blots were imaged using a Li-Cor Odyssey infrared imaging system (Li-Cor), and signals were quantified using Emperia software (Li-Cor). Results were normalized to the mean of each corresponding WT replicates set and represented as a fold change relative to WT.

### RT-qPCR

Total RNA was purified from brain homogenates using RNAeasy Kit (Qiagen) as previously described^37^. The threshold-crossing value was normalized to internal control transcripts (*18S*, *Actb*, and *Pgk1*). Results were normalized to the mean of each corresponding WT replicate set and represented as a fold change relative to WT.

### Statistics

Statistical analyses were performed with Prism 9 (Graph Pad) unless specified otherwise. D’Agostino & Pearson, Shapiro–Wilk, and Kolmogorov-Smirnov normality tests were performed to verify normality. For data that fulfilled normality requirements, unpaired *t* tests were used for pairwise comparisons of WT and OGT^C921Y^ data or Two-way Anova for multiple comparisons were used. For data sets that did not fulfil normality, Mann-Whitney tests were used for pairwise comparisons.

## Results

### OGT^C921Y^ mice display postnatal growth development delay and hyperactivity

In patients, OGT-ID variants associate with balance difficulties, ataxia and hypotonia, suggesting possible locomotor defects in addition to ID and behavioural deficits^63^. To investigate whether the OGT^C921Y^ variant phenocopies such deficits in mice, we first assessed a range of behavioural traits, including locomotion, anxiety, compulsivity, learning and memory in a mouse line carrying this variant, the generation of which has recently been reported^37^. As part of our overall behavioural screening, animal body weight was monitored weekly. During the initial period after weaning (3 to 8 weeks old), no difference in body weight was observed between OGT^WT^ and OGT^C921Y^ mice suggesting similar developmental trajectories between both genotypes. However, we observed statistically significant lower gain in body weight in the OGT^C921Y^ mice from 9 to 20 weeks old compared to the WT mice (Time x Genotype *p* = <0.0001, 2-way ANOVA) (**Fig. 1A**), suggesting that OGT^C921Y^ mice show postnatal growth development delay.

**Figure 1:**
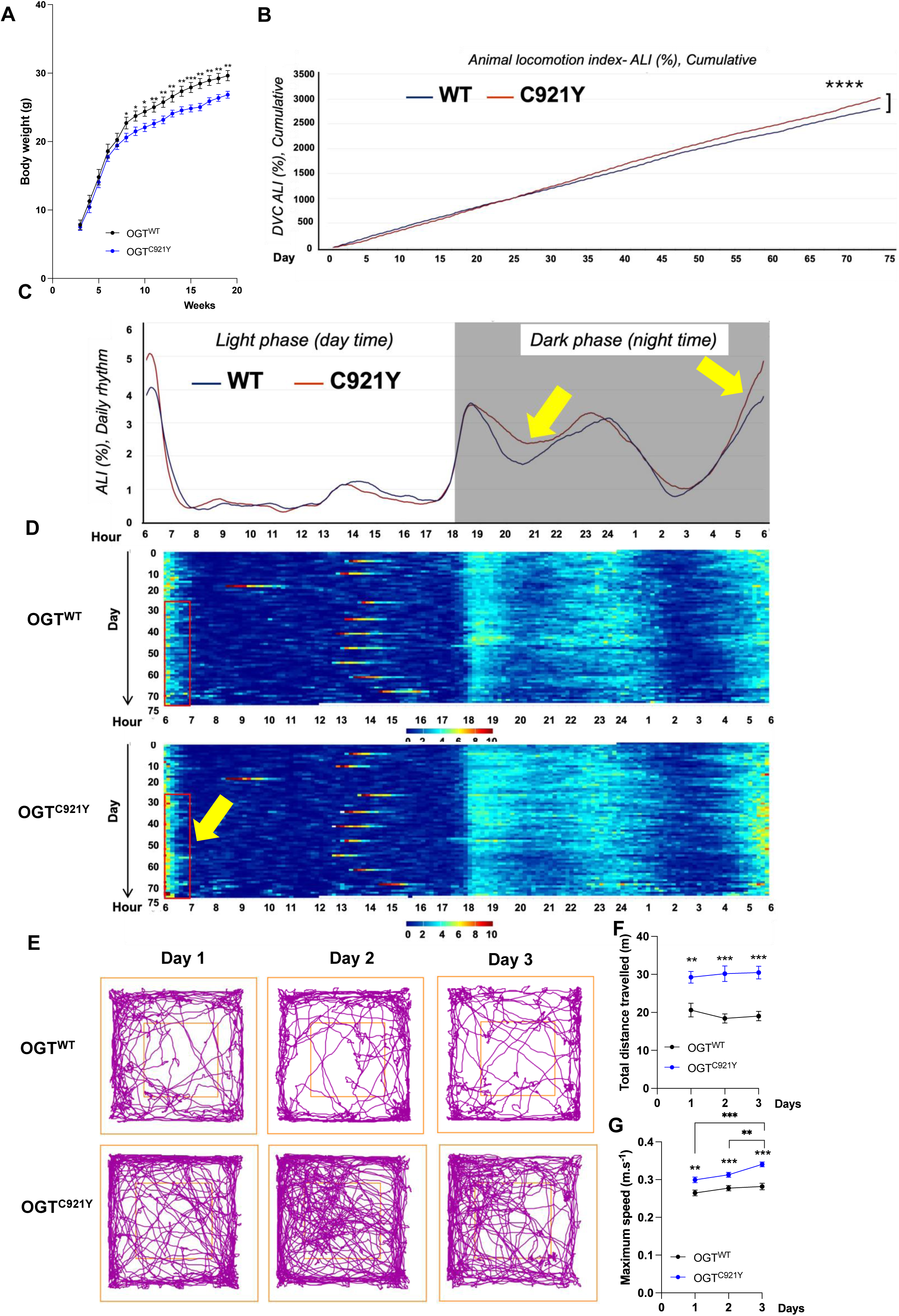
OGT^C921Y^ mice show postnatal growth development delay and increased spontaneous activity. Significance is shown as \**p* < 0.05, \*\**p* < 0.01, \*\*\**p* < 0.001 and ****p < 0.0001. (A) Body weight of male OGT^WT^ (n=15) and OGT^C921Y^ (n=16) mice from 3 weeks to 20 weeks old. 2-way ANOVA (Alpha, 0.05) followed by Tukey’s column comparisons. (B) Line chart representing the cumulative locomotion index (x-axis, days) for 75 days. Two-way ANOVA (Alpha, 0.05) followed by Tukey’s column comparisons (OGT^WT^, n = 10 housed in 5 cages; OGT^C921Y^, n = 10 housed in 5 cages). (C) Line chart representing the cumulative locomotion index with light (6 am – 6 pm) and dark (6 pm – 6 am) periods in DVC cages over a longitudinal period up to 75 days (x-axis showing time in hours, 24-h format). The arrows indicate increased nocturnal activity by OGT^C921Y^ mice compared to WT littermates. (D) Heat maps representation of the cumulative locomotion index with light and dark periods in DVC cages for 75 days (x-axis showing time in hours over 24-hour period each day, y-axis showing days with Day 31 as the starting point on top). The arrow indicates increased nocturnal activity by OGT^C921Y^ mice compared to WT littermates. (E) Representative tracking plot over three consecutive days of a male OGT^WT^ and male OGT^C921Y^ mice in the open field arena. (F) Distance travelled over three consecutive days of male OGT^WT^ (n = 15) and male OGT^C921Y^ (n = 16) mice in the open field arena. 2-way ANOVA (Alpha, 0.05) followed by Tukey’s column comparisons. (G) Maximal speed displayed by male OGT^WT^ (n = 15) and male OGT^C921Y^ (n = 16) mice in the open field arena. 2-way ANOVA (Alpha, 0.05) followed by Tukey’s column comparisons.

Shortly after weaning, we placed the animals in digitally ventilated cages to continuously monitor the patterns of spontaneous activity in a non-invasive manner^38^. For this purpose, we tracked animal activity longitudinally, starting at postnatal day 30 until 105 days of age (data collected over 75 consecutive days; n = 10 per group, housed in five separate cages). These experiments revealed hyperactive patterns of home cage activity by the OGT^C921Y^ mice, compared to the WT littermates, as early as day 35 (*p* = <0.0001, 2-way ANOVA) (**Fig. 1B**). Intriguingly, this period (postnatal 9-20 weeks) also marks the stage reflecting the observed postnatal growth development delay in the OGT^C921Y^ mice (**Fig. 1A**). Evaluation of daily rhythms and home cage activity during light/dark periods revealed that the OGT^C921Y^ mice exhibited frequent periods of hyperactivity during the dark phase (night time, active time for rodents), compared to the WT (**Fig. 1C, D**). However, the resting time (light phase, day time) was similar between the genotypes (**Fig. 1C, D**) ruling out potential disturbances in circadian rhythms, which otherwise could affect their normal day/night patterns of home cage activity.

We next investigated general locomotor activity in an open field arena. Mice were placed in open arenas and allowed to explore for 10 min on three consecutive days to evaluate both exploration of, and habituation to, an unfamiliar environment. Over the total three day period, OGT^C921Y^ mice showed an increase in activity in the arena as reflected by an increase in total distance travelled (Day 1 *p* = 0.0011; Day 2 *p* = <0.0001; Day 3 *p* = <0.0001, 2-way ANOVA) and maximum speed (Day 1 *p* = 0.0013; Day 2 *p* = 0.001; Day 3 *p* = <0.0001, 2-way ANOVA) compared to WT mice (Fig. **1E, F, G**). To assess whether this hyperactivity was due to higher anxiety levels in OGT^C921Y^ mice, we evaluated the fraction of time spent in the periphery versus centre of the arena. Over the three days of testing, OGT^C921Y^ mice demonstrated thigmotaxic behaviour and increased distance travelled in both periphery (Day 1 *p* = 0.0004; Day 2 *p* = <0.0001; Day 3 *p* = <0.0001, 2-way ANOVA) and centre areas (Day 1 *p* = 0.014; Day 2 *p* = 0.0012; Day 3 *p* = 0.0193, 2-way ANOVA), but no preference for the periphery of the arena compared to WT, as shown by similar time spent in the periphery (Day 1 *p* = 0.1771; Day 2 *p* = 0.2657; Day 3 *p* = 0.8885, 2-way ANOVA) and centre areas (Day 1 *p* = 0.1785; Day 2 *p* = 0.2543; Day 3 *p* = 0.8849, 2-way ANOVA) between both genotypes (**Fig. S1A-D**). During Elevated Plus Maze (EPM), a test to assess elevated and open space anxiety, OGT^C921Y^ mice covered more distance (*p* = <0,0001, *t*-test) and performed higher number of entries in the centre (*p* = <0,0001, *t*-test) resulting in an increase in open/close time ratio (*p* = 0,0063, *t*-test) (**Fig. S1E-H**). Similarly, OGT^C921Y^ mice travelled more distance (0-5 min *p* = 0.009; 5-10 min *p* = <0.0001, 2-way ANOVA) and displayed frequent light-dark transitions compared to WT mice during the dark-light paradigm test (0-5 min *p* = 0.001; 5-10 min *p* = 0.3374, 2-way ANOVA), with no difference in time spent in the light compartment between both genotypes (0-5 min *p* = 0.0926; 5-10 min *p* = 0.0608, 2-way ANOVA) (**Fig. S1I-L**). These findings potentially rule out anxiety-like behaviour in both the OGT^C921Y^ and the WT mice.

We also assessed motor skills, including sensorimotor coordination and balance. No differences were observed in mean speed (*p* = 0.9056, *t*-test) or time spent on the rotarod (*p* = 0.814, *t*-test) between WT and OGT^C921Y^ mice (**Fig. S2A, B**). In the pole test, both OGT^C921Y^ and WT mice were able to perform a t-turn and reach the ground with similar scores (*p* = 0.1523, Mann-Whitney test) (**Fig. S2C**). During static rod tests, OGT^C921Y^ mice spent more time than their WT littermates to complete the required t-turn (35 mm *p* = 0.0018; 28 mm *p* = 0.0003; 22 mm *p* = 0.1201; 15 mm *p* = 0.0005; 10 mm *p* = <0.0001, 2-way ANOVA) although all mice reached the goal platform (**Fig. 2A, B, C**). Taken together these data rule out gross defects in motor coordination and balance in the OGT^C921Y^ animals, while revealing a degree of hyperactivity and stereotypy.

**Figure 2:**
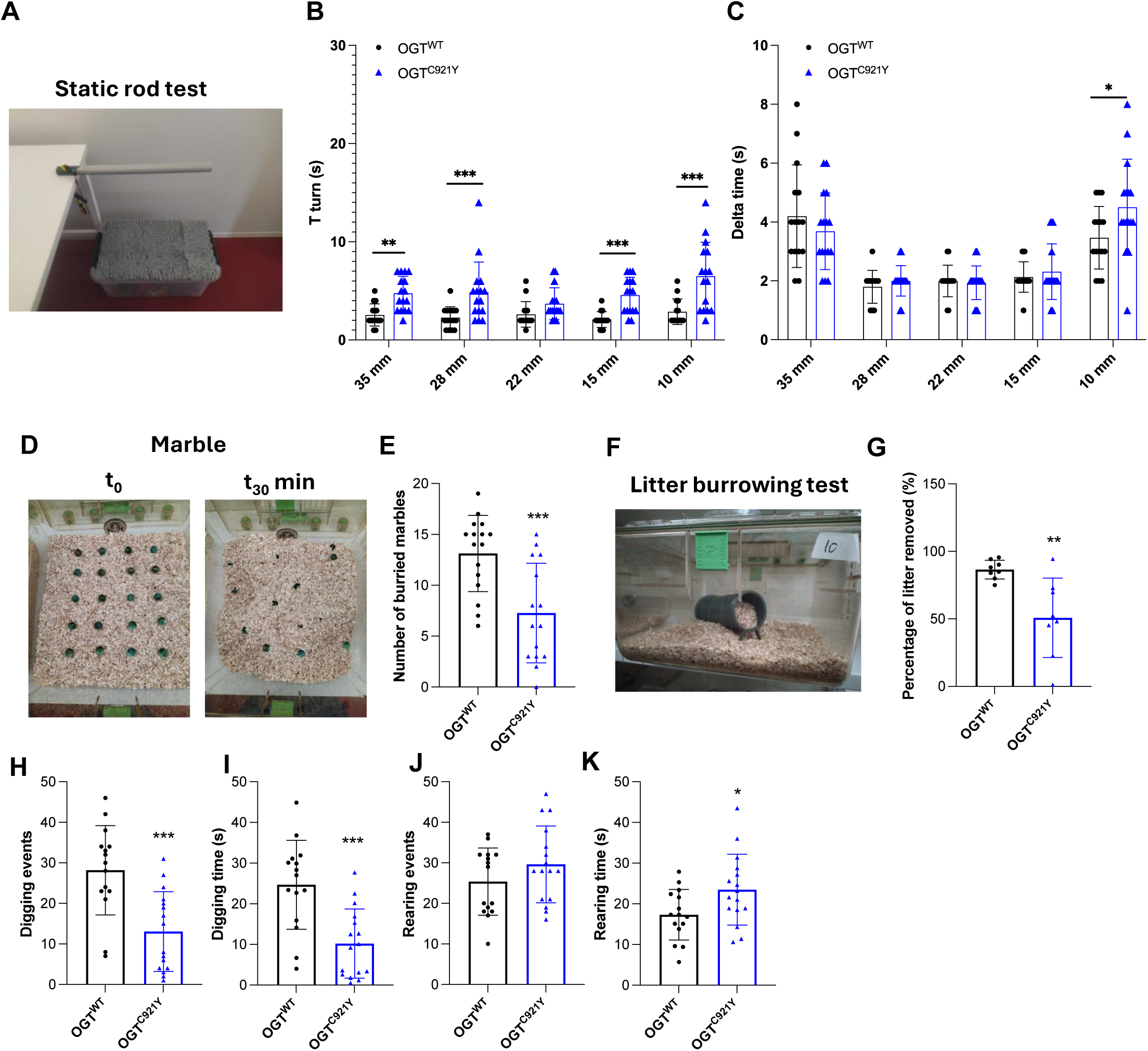
Motor, digging and burying activity. Significance is shown as * *p* < 0.05, ** *p* < 0.01 and *** *p* < 0.001. Student *t* test was used for statistics. (A) Representative image of experimental set-up for the static rods test. (B) Time to t-turn of male OGT^WT^ (n = 15) and OGT^C921Y^ (n = 16) during the static rods test. (C) Delta time defined as (Total time – time to turn) of male OGT^WT^ (n = 15) and OGT^C921Y^ (n = 16) during the static rods test. (D) Representative images of the marble test at 0 min (T0) and 30 min (T30). (E) Number of buried marbles by male OGT^WT^ (n = 15) and OGT^C921Y^ (n = 16) during the marble test. (F) Representative image of the litter burrowing test. (G) Percentage of litter removed by male OGT^WT^ (n = 15) and OGT^C921Y^ (n = 16) during the litter burrowing test. (H) Number of digging events by male OGT^WT^ (n = 15) and OGT^C921Y^ (n = 16) during 3 min observation. (I) Time spent digging by male OGT^WT^ (n = 15) and OGT^C921Y^ (n = 16) during 3 min observation. (J) Number of rearing events by male OGT^WT^ (n = 15) and OGT^C921Y^ (n = 16) during 3 min observation. (K) Time spent rearing by male OGT^WT^ (n = 15) and OGT^C921Y^ (n = 16) during 3 min observation.

### OGT^C921Y^ mice exhibit reduced digging behaviour

In order to further characterize the stereotypic phenotypes that could reflect ASD features observed in some OGT-ID patients^63^, we assessed compulsive and repetitive behaviour. The OGT^C921Y^ showed a reduced number of buried marbles in a marble burying test (*p* = 0.0002, *t*-test) (**Fig. 2D, E**) and a reduction in litter displacement in a burrowing test (*p* = 0.0107, *t*-test) (**Fig. 2F, G**) compared to WT mice. To investigate whether the reduction in burying/burrowing activity was due to reduced digging behaviour, WT and OGT^C921Y^ were individually placed in cages containing 5 cm deep litter. The number of digging events and time spent digging during 3 min were measured. The OGT^C921Y^ mice exhibit both reduced number of digging events (*p* = 0.0004, *t*-test) and digging time (*p* = 0.0003, *t*-test) compared to WT mice (**Fig. 2H, I**). These findings were not due to lower activity of the OGT^C921Y^ mice, but rather to an increase in rearing events (*p* = 0.1978, *t*-test) and time spent rearing (*p* = 0.0317, *t*-test) compared to WT mice (**Fig. 2J, K**). The number of self-grooming events (*p* = 0.7421, *t*-test) and time spent self-grooming (*p* = 0.3785, *t*-test) were similar between the two genotypes (**Fig. S2D, E**). In addition, OGT^C921Y^ mice removed the same amount of material as WT mice during the nesting test (*p* = 0.9774, *t*-test) (**Fig. S2F**).

### OGT^C921Y^ mice manifest features of impulsive behaviour and enhanced long term memory recall

We next investigated the effect of the OGT^C921Y^ variant on memory. First, we evaluated spontaneous alternation using a T-maze test as a measure of working spatial memory. The mice were placed in a T-shape maze and allowed to choose freely between right (R) and left (L) arms for 7 successive trials (**Fig. 3A**). Both OGT^C921Y^ and WT mice show similar percentage of spontaneous alternation (*p* = 0.2347, *t*-test) (**Fig. 3B**), suggesting intact short-term spatial working memory in the OGT^C921Y^ mice. Nevertheless, the OGT^C921Y^ mice display a significantly reduced latency of choice (*p* = 0.0175, Mann-Whitney test) compared to WT mice (**Fig. 3C**), suggesting possible impulsive behaviour.

**Figure 3:**
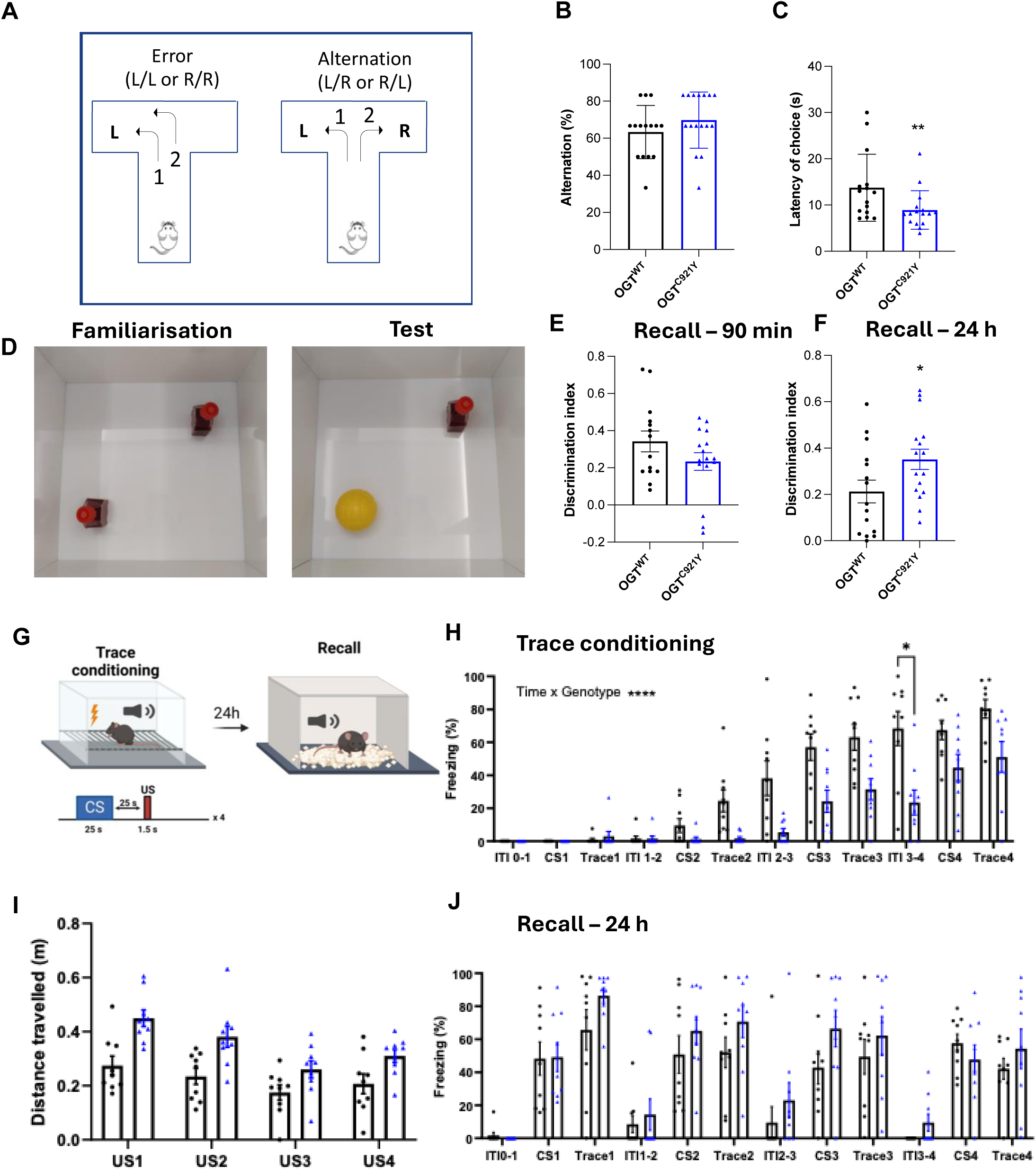
OGT^C921Y^ show impulsivity, enhanced long term memory and delayed aversive associated learning. Significance is shown as \**p* < 0.05, \*\**p* < 0.01 and \*\*\**p* < 0.001. (A) Schematic of the T-maze paradigm. Mice are free to explore both arms for 7 additional trials. Each choice and latency were recorded. (B) Percentage of correct alternation (L-R/R-L sequences) of OGT^WT^ (n = 15) and OGT^C921Y^ (n = 16) during the T-maze test. Student *t* test was used for statistics. (C) Latency of arm entry of OGT^WT^ (n = 15) and OGT^C921Y^ (n = 16) during the T-maze test. Student *t* test was used for statistics. (D) Representative image of the Novel Object Recognition (NOR) test. During the familiarization phase, mice are free to explore an arena with two identical objects. During the test phase, mice were free to explore the same arena where one of the familiar objects has been replaced by a novel object to assess short (90 min) and long (24 h) memory. (E) Discrimination index of OGT^WT^ (n = 15) and OGT^C921Y^ (n = 16) during the short term (90 min) NOR test. Student *t* test was used for statistics. (F) Discrimination index of OGT^WT^ (n = 15) and OGT^C921Y^ (n = 16) during the long term (24h) NOR test. Student *t* test was used for statistics. (G) Diagram showing the behavioural paradigm (top), tone, conditioned stimulus (CS), trace period, and unconditioned stimulus, foot shock (US) lengths (bottom). (H) Freezing behaviour of OGT^WT^ and OGT^C921Y^ (n = 9 per group) during the acquisition session of the aversive conditioning. ITI, Intertrial interval. 2-way ANOVA time x genotype interaction *p* < 0.0001, F(11,176) = 5.572. (I) Locomotor activity of OGT^WT^ and OGT^C921Y^ (n = 9 per group) evoked by the US. The data represent the 3 s bin following the foot shock. Two-way ANOVA time x genotype interaction *p* = 0.3323, F(3,48) = 1.166. (J) Freezing behaviour of OGT^WT^ and OGT^C921Y^ (n = 9 per group) in the recall session where the CS were presented without US. The trace period refers to the 25 s following the CS presentation. Two-way ANOVA time x genotype interaction *p* = 0.5337, F(11,176) = 0.9082.

We also investigated novelty-associated memory using the Novel Object Recognition (NOR) paradigm. We used two different sets of objects to assess both short (90 min) and long (24 h) term memory. During the familiarisation phase, the mice were placed in an open field arena with two identical objects for 10 min. After 90 min or 24 h, mice were placed in the same arena containing one familiar object and one novel object (**Fig. 3D**). Time spent exploring each object was recorded allowing 20 s total exploration of both objects. Discrimination indexes were approached by calculating (t_novel_ – t_familiar_) / t_total_. After a 90 min interval, OGT^C921Y^ mice showed a similar discrimination index as WT mice (*p* = 0.1482, *t*-test), suggesting normal short term memory function in this assay (**Fig. 3E**). However, while both WT and OGT^C921Y^ show discrimination indices greater than 0.2 after 24 h, OGT^C921Y^ mice showed significantly greater discrimination index compared to WT mice (*p* = 0.0434, *t*-test), potentially indicating enhanced long term memory recall (**Fig. 3F**).

### OGT^C921Y^ mice exhibit impaired plasticity during associative learning

To assess learning and memory function, we subjected OGT^C921Y^ mice and WT littermates to aversive Pavlovian trace conditioning - an established hippocampus- and cortex-dependent learning and memory paradigm where mice learn to associate a neutral tone with an aversive foot shock in a non-contiguous manner. Learning involves the presentation of a tone as the conditioned stimulus (CS), followed by a trace period, after which a foot shock is delivered as the unconditioned stimulus (US) (**Fig. 3G**). We used freezing, defined as the complete cessation of movement (except for breathing), as the conditioned response (CR) and a surrogate measure of short-term adaptation and aversive learning. Freezing behaviour was monitored during the acquisition phase and again 24 h later during a recall session (**Fig. 3G**). We observed that OGT^C921Y^ mice exhibited significant learning deficits, as indicated by a delay in freezing behaviour compared to WT littermates (Time x Genotype *p* = <0.0001, 2-way ANOVA) (**Fig. 3H**). This learning deficit cannot be attributed to somatosensory anomalies, as both genotypes displayed comparable responses to the US, as indicated by locomotor activity evoked by the foot shock (Time x Genotype *p* = 0.3323, 2-way ANOVA) (**Fig. 3I**). Furthermore, mutant mice reached the same CR levels as WT mice by the fourth CS-tone presentation (**Fig. 3H),** suggesting unimpaired auditory function. During the 24 h recall, there were no differences in freezing behaviour in OGT^C921Y^ mice compared to WT mice (Time x Genotype *p* = 0.5337, 2-way ANOVA) (**Fig. 3J**), suggesting an impaired plasticity during the acquisition phase. Taken together, these data suggest that OGT^C921Y^ exhibit impaired plasticity during associative learning.

### OGT^C921Y^ mice exhibit reduced skull size and shape deformation

OGT-ID is often associated with microcephaly or craniofacial deformities, which were also recapitulated in our initial report using a small cohort (n < 5) of 2 month old OGT^C921Y^ mice^37^. Here we aimed for a comprehensive skull shape analyses using micro-computed tomography (μCT) in larger cohorts of older (5 month) mice (WT, n = 14; OGT^C921Y^, n = 16). Average skull shapes were superimposed for OGT^WT^ and OGT^C921Y^ mice and skull shape differences between the two groups were analysed. The OGT^C921Y^ skulls were smaller in both the anterior and posterior areas, whereas the top of the skull appeared more curved compared to the WT skulls (**Fig. 4A**). We performed similar analyses on the previously reported μCT data collected from 2 month old animals^37^ to detect possible temporal aspects to these defects. The 2 month old OGT^C921Y^ mice show similar but more prominent shape differences than the 5 month old OGT^C921Y^ mice, suggesting an age-dependent convergence of skull shapes (**Fig. 4A**). Next, surface landmarks were used to perform Principal Component Analysis (PCA) allowing comparison of skull shape independently of size differences **(Fig. S3B)**. Skulls from 5 month old OGT^WT^ and OGT^C921Y^ mice were separated along the first principal component (PC1) with a higher dispersion in the OGT^C921Y^ group suggesting heterogeneity in skull shape deformation phenotype penetrance. More than 63 % of the OGT^C921Y^ skulls display a positive PC1 value while 93 % of the WT skulls display a negative PC1 value. The skull of the OGT^C921Y^ mice has a shorter and rounder shape than the average WT mice (**Fig. 4B**). We performed Euclidean Distance Matrix Analysis (EDMA) using 45 relevant landmarks from the Richtsmeier laboratory resource (https://getahead.la.psu.edu) to identify the key regions contributing to the shape differences (**Fig. S3A**). At 5 months of age, 55% of the distances were at least 2% shorter in the OGT^C921Y^ skulls than in WT skulls, with the largest differences affecting the cranial base length (**Fig. S3C**). Similarly, in skulls from 2 month old animals, distances that are shorter than 5% in the OGT^C921Y^ skulls were predominantly located at the base of the skull (**Fig. S3C**). The main bone contributing to the cranial base length is the sphenoid, and shortening in the sphenoid bone determines the curvature of the cranial vault postnatally^64^. This may indicate that defects in the postnatal development of the skull could cause the observed phenotype in the OGT^C921Y^ mice. Similar skull shape deformities have been reported in mouse models of Fragile X syndrome^65^ and skeletal defect syndromes, such as osteogenesis imperfecta^66^.

**Figure 4:**
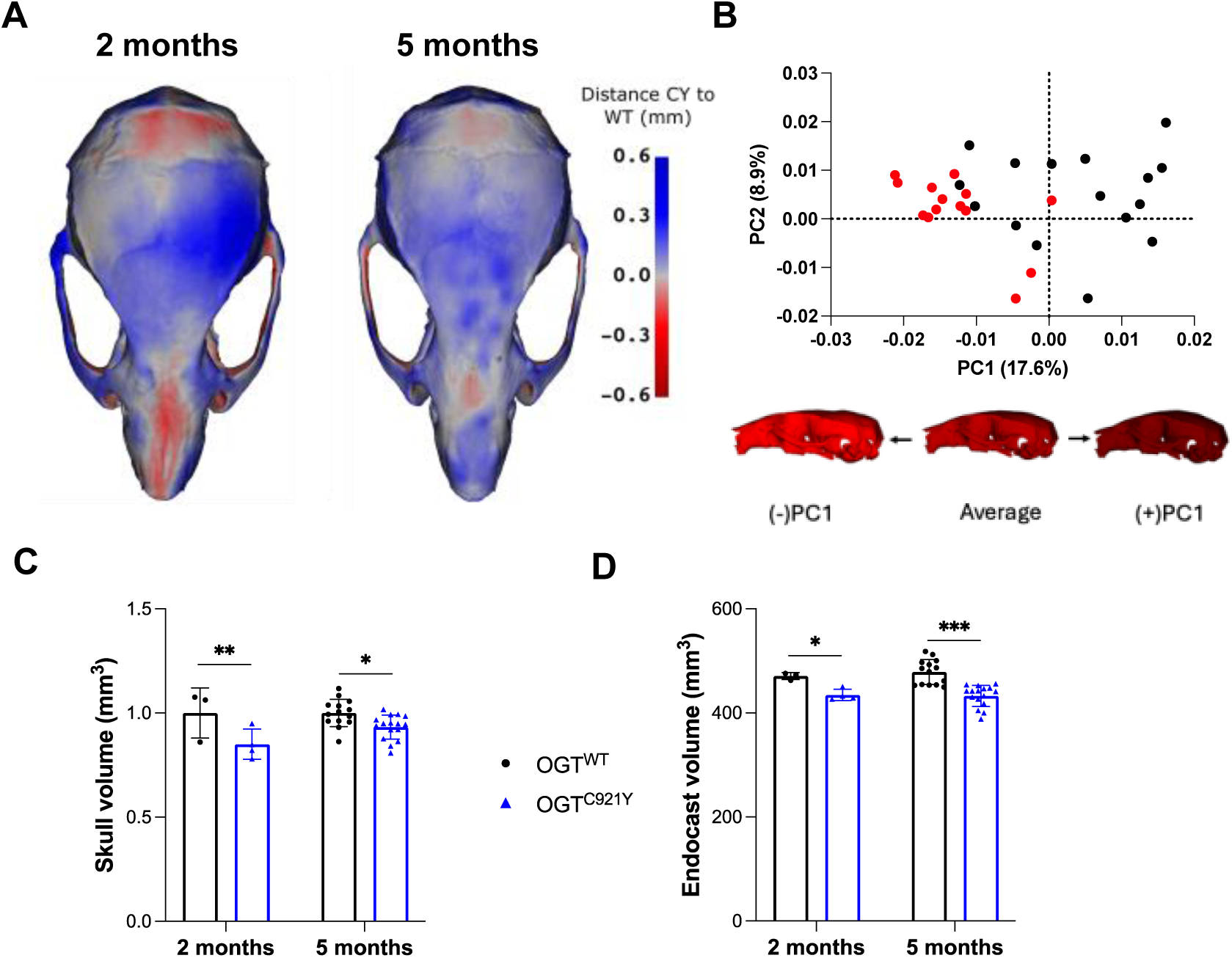
Micro-computed tomography of OGT-ID mouse skulls indicates a reduced endocast volume and shorter skull length. Significance is shown as * *p* < 0.05, ** *p* < 0.01 and *** *p* < 0.001. (A) Heatmaps obtained of the average OGT^C921Y^ skull (n = 16) to the average OGT^WT^ skull (n = 14). Red and blue regions indicate that the average OGT^C921Y^ skull is smaller or larger respectively than the average OGT^WT^ in those areas. (B) PCA biplot of PC1 and PC2 of the surface landmarks of 20 weeks old male OGT^WT^ and OGT^C921Y^ skulls (top). Percentages in the axis indicate the explained variance. Deformations in PC1 showing shape differences across this axis are shown (bottom). (C) Volume of the skulls of 8 weeks and 20 weeks old male OGT^WT^ and OGT^C921Y^ mice. Student *t* test was used for statistics. (D) Volume of the extracted endocranial cavity of 8 weeks and 20 weeks old male OGT^WT^ and OGT^C921Y^ mice. Student *t* test was used for statistics.

At both ages (i.e., 2 and 5 months old), the OGT^C921Y^mice, the reduction in cranial base distances and overall reduction in skull size were obvious (2 months *p* = 0.0016; 5 months *p* = 0.0101, 2-way ANOVA) (**Fig. 4C**). From 3D reconstruction of the skulls, we determined the endocranial cavity volume as an indirect measure of the brain volume. At both ages tested, OGT^C921Y^ mice had a smaller internal cranial volume compared to their WT littermates (2 months *p* = 0.013; 5 months *p* = <0.0001, 2-way ANOVA) (**Fig. 4D**), suggesting a smaller brain volume. Taken together, these findings indicate that the OGT^C921Y^ mice exhibit antero-posterior skull growth defects leading to reduced skull size and shape deformation, suggesting dysmorphic features and reduced brain size.

### OGT^C921Y^ display reduced cortical thickness and hypoplastic changes in brain structures

Neocortex hypoplasia, changes in cortical thickness and white matter integrity defects have been reported in mouse models of neurodevelopment disorders including ASD^67^, CHARGE^68^ and Rett^69^ syndromes. To assess whether the reduction in brain size is general or localized to specific brain regions, we employed magnetic resonance imaging (MRI). Specifically, we performed volumetry analysis of anatomically well-defined brain regions based on high-resolution MRIs of 50 µm isotropic resolution, covering both the grey and white matter. The OGT^C921Y^ mice showed a reduced total brain volume compared to WT (**Fig. 5A**). Regional volumetric analyses showed a reduction in absolute volumes in most brain regions (**Fig. S4**). When expressed as regional relative volume (RRV) variation, the OGT^C921Y^ mice showed significantly reduced (group median) volumes of several brain regions including the frontal lobe of the cerebral cortex, corpus callosum, basal forebrain, globus pallidus, internal capsule and stria terminalis (**Fig. 5C, S5**). In contrast, RRV was increased in the lateral septum, medulla, hippocampus and hypothalamus regions of the OGT^C921Y^ mice compared to WT mice (**Fig. 5C, S5**). Cortical thickness analysis revealed significantly thinner cortex bilaterally in posterior regions in the OGT^C921Y^ mice compared to WT mice, with differences on the order of 100 µm (**Fig. 5B**).

**Figure 5:**
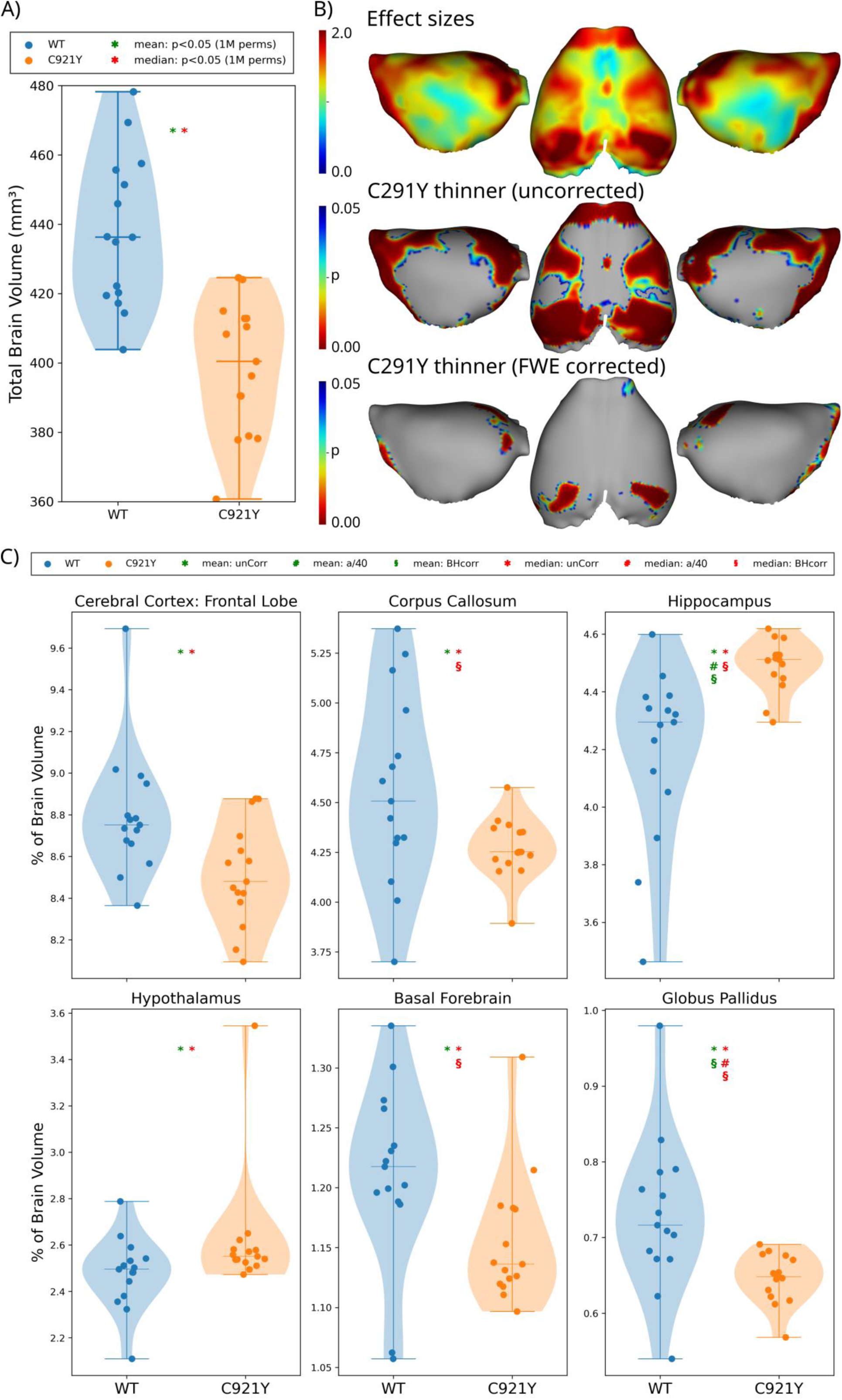
OGT^C921Y^ mice show changes in regional brain volume and reduced cortical thickness. Volumetry and cortical thickness from high resolution T1-weighted Magnetic Resonance (MR) images. (A) Total brain volume of 20 weeks old male OGT^WT^ and OGT^C921Y^ mice. Groupwise total brain volume where each dot represents one subject. The horizontal lines correspond to group extrema and median. Asterisks (*) mark significance (p < 0.05) based on permutation tests (1M permutations) of either the mean (green asterisk) or median (red asterisk) of the two groups. (B) Heatmaps of group differences in cortical thickness between male OGT^WT^ and OGT^C921Y^ mice. Maps are effect sizes (top row), p-values of significant differences both uncorrected for multiple comparisons (middle row) and corrected with family-wise error (FWE). N = 15 per genotype. One mutant was not perfused correctly and removed from analysis. (C) Regional brain volumes represented as percentage of whole brain volume of male OGT^WT^ and OGT^C921Y^ mice. Asterisks (*) indicate uncorrected significance (p < 0.05), based on permutation tests (100k permutations for each region). Pound symbols (#) indicate significance below alpha (0.05) divided by total number regions tested (40) for mean (green) and median (red), respectively. Section signs (§) indicate significance (p < 0.05) with p-values adjusted for false discovery rate (Benjamini-Hochsberg, BH). Y-axes are scaled to individual ROIs to highlight group variation and difference.

We next assessed potential local brain microstructural defects using diffusion kurtosis imaging (DKI)^70–73^, which provides sensitive indices of water mobility (mean diffusivity, MD) and diffusion directionality (fractional anisotropy, FA) in tissue as well as markers of tissue complexity (mean kurtosis, MK). Collectively, the MRI markers employed here are known to be sensitive to subtle tissue alterations in both rodent^41,42,48,74–76^ and human brain^77–80^. We observed no difference in these DKI metrics between OGT^C921Y^ and WT mice in any of the 20 automatically segmented regions including neocortex (**Figs. S6-9**). Taking together, these MRI data suggest that OGT^C921Y^ show preserved brain microstructure at the resolution (150 µm x 150 µm x 250 µm) of the DKI experiments yet reduced cortical thickness.

### OGT^C921Y^ mice exhibit dysmorphic features in superficial cortical layers, predominantly affecting the cingulate

Previous studies indicate that congenital dysplasia in cortical organization manifests as a range of malformations including disrupted cortical laminar organization, neuronal heterotopia in the subcortical white matter, misplaced neurons in cortical lamina I, clustering of neurons in the grey matter, and the presence of dysmorphic neurons^81–83^. To evaluate cortical cytoarchitecture at the microscopic level and for detecting any features suggestive of focal cortial dysplasia (FCD) in OGT^C921Y^ mice, we subjected the brain sections to histological analyses using H&E (for cells), cresyl violet (for neurons), luxol fast (for white matter) staining (**Fig. S10**). In the H&E analyses, we observed an overall normal 6-layered cortical organization across the regions involved in primary sensorimotor modalities, including the somatomotor cortex M1/M2, primary somatosensory cortex S1/S2, auditory cortex and primary visual cortex V1 (**Fig. S11**). Intriguingly, part of the cingulate (retrosplenial Area 29/Area 30, according to Paxinos and Franklin) found within the paramedian portion (sagittal section, interaural 0.36) showed FCD in 5 out of 6 animals in the OGT^C921Y^ cohort (**Fig. 6A-C**, compared to WT). The most conspicuous microscopic finding suggestive of focal cortical dysplasia was seen in the form of pseudosulcus formation in the cingulate, such that the superficial cortical layers (I-III) appeared to be displaced inwards (**Fig. 6B-C**, black arrow), and resembled polymicrogyria similar to those observed in other congenital brain malformations^84,85^. In two extreme cases, nodular arrangements of cells within the superficial layers (II/III bordering IV), with a central halo containing loosely arranged eosinophilic tissue, were observed (**Fig. 6B-C**, yellow arrow). Having ruled out gross structural alterations (except in the cingulate cortex), we next assessed cortical cell density, which would reveal hypo or hyperproliferative neurodevelopmental anomalies. Total cell density analyses in the neocortical regions (H&E stained serial sections, reflecting both neuronal and non-neuronal cells), did not indicate drastic differences between the two groups in the major cortical regions examined (**Fig. 6D**).Staining analyses using cresyl violet indicated that the nodular collections in the layers II/III bordering IV were predominantly neuronal cell bodies (Fig. 6E). In luxol fast stained sections, these malformations contained collections of ectopic white matter, which populated the center of nodular malformations (**Fig. 6F**) and/or found as patchy deposits in the vicinity of superficial layers (II-IV) (**Fig. 6C**).To further establish the cell identity in the cortical malformations, we performed immunofluorescence analyses using neuronal nuclei marker (NeuN) and astroglial marker glial acidic fibrillary protein (GFAP). These analyse further corroborated the notion that the foci of FDC in the cingulate of OGT^C921Y^ mice were predominantly populated by neuronal cells and were devoid of astrocytes, which otherwise would indicate reactive astrogliosis seen in states of brain trauma (**Fig. 6G-H**, compare WT in Fig. 6G to OGT^C921Y^ in Fig. 6H). Furthermore, normal localization of astroglial cells in the subcortical white matter was evident in the WT and OGT^C921Y^ cohorts (**Fig. 6G-H**, merged images in the panels on right). In parallel, multiplex IHC analyses confirmed that the nodular cortical malformations predominantly affected layers II-V, as indicated by the distribution of neuronal markers Reelin and POU class 3 homeobox 2 (POU3F2), and were populated by neurons and oligodendrocytes in the vicinity (**Fig. S12**). Taken in conjunction with the MRI (reduced cortical thickness), the cortical dysplasia in revealed by the histology analyses suggest a neurodevelopmental component to OGT-ID affecting the cortical superficial layers, which in OGT^C921Y^ mice predominantly affects the cingulate.

**Figure 6:**
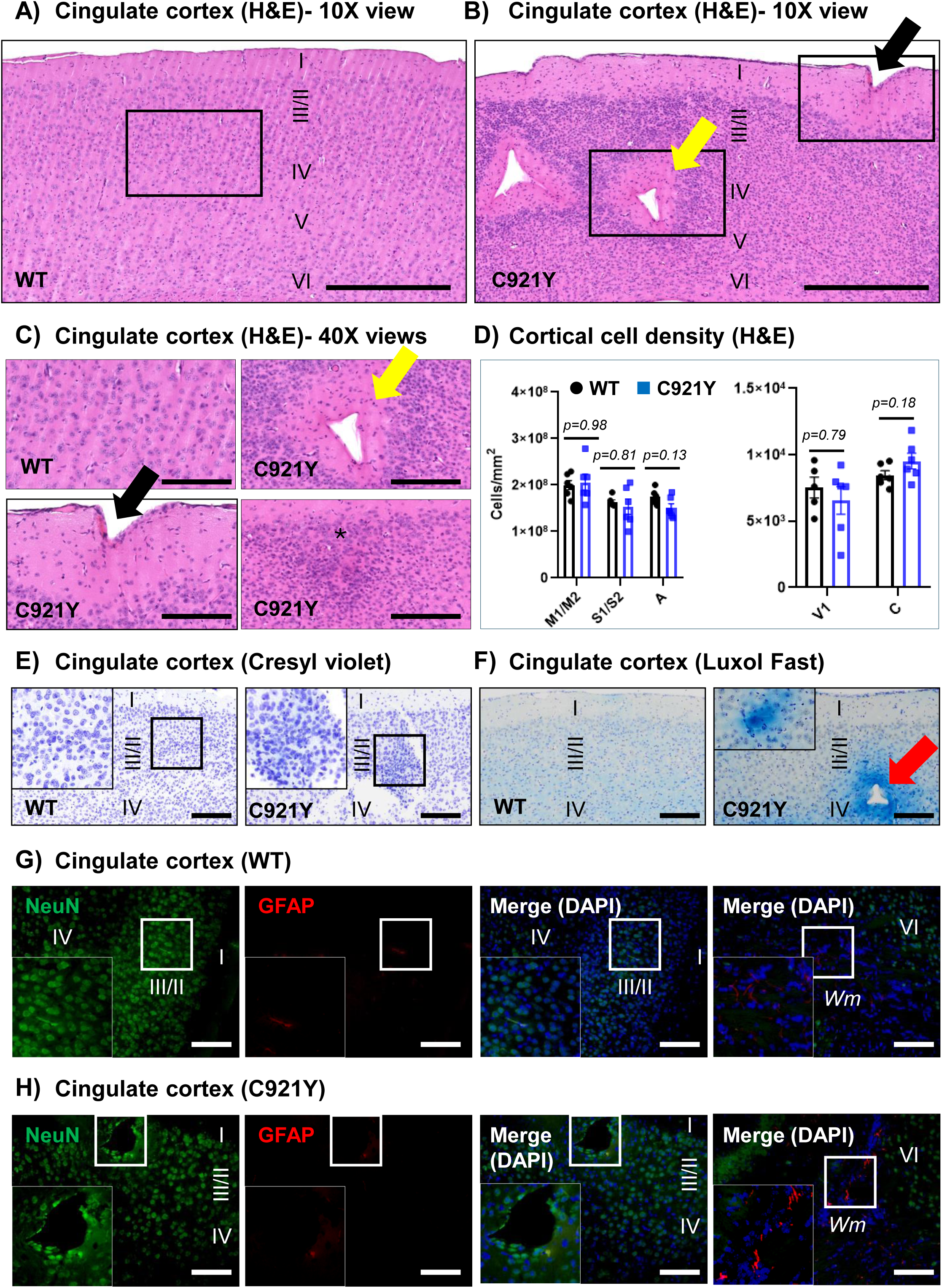
Pseudosulcus formation, cortical dysplasia and ectopic white matter in the cingulate cortex of OGT^C921Y^ mice. (A-B) Representative low magnification (10X) sagittal views of WT (in A) and OGT^C921Y^ mice (in B), showing cortical malformations in the cingulate cortex of the latter by H&E staining. The black arrow points to instances of pseudosulcus formation and the yellow arrow points to instances of cortical dysplasia observed in the layers II-IV (scale bar = 500 µm). (C) Representative high magnification (40X) views extracted from the insets in A except one marked with *, which is depicting nodular arrangement of cells in a brain section from a different OGT^C921Y^ animal (scale bar = 100 µm). (D) Regional cell density assessed in 10X views (H&E staining) of two-serial sections from WT and OGT^C921Y^ mice. Error bars indicate Mean ± SEM (n = 6 / group; Mann-Whitney test, none significant). Regions examined include: (coronal sections) primary motor cortex M1/M2, primary somatosensory cortex S1/S2, auditory cortex (A); (sagittal sections) visual cortex V1 and cingulate cortex (C). Also see Fig. S10 depicting coordinates and region demarcations (dashed rectangles) used in the cell density analyses. (E-F) Representative low magnification (10X) sagittal views showing cresyl violet staining for neuronal cell bodies (in E) and myelin staining by luxol fast (in F). Notice the instances of ectopic white matter (red arrows) in the cingulate cortex of OGT^C921Y^ mice (scale bar = 200 µm). in 6F.

### Quantitative proteomics analyses revealed distinctly perturbed molecular pathways in prefrontal cortex of the OGT^C921Y^ mice

In order to decipher the molecular phenotypes that could be associated with the cortical defects in the OGT-ID mutant strain, we performed proteome analysis on the prefrontal cortex (PFC) of male OGT^C921Y^ and WT mice using label-free quantification. A total of 5488 proteins were identified with a 1 % false discovery rate which we then ranked as top 50 most affected (and potentially revealing) factors. This ranking was based on defined cut-off criteria for both the upregulated (*p* value ≤ 0.05; log-2 fold change ≥ 1.45) and down-regulated proteins (*p* value ≤ 0.05, log-2 fold change ≤ −1.33). Then we performed pathways enrichment and downstream analyses using the built-in tools of the STRING database (**Fig. 7A, B**). To begin with, the expression of OGA was significantly downregulated in the cortex of OGT^C921Y^ (**Fig. 7B**) in the mass spectrometry analyses, as well as in western blotting analyses of the cortex (*p* = 0.0223, *t*-test) (**Fig. S13A, C**) as we have reported previously^60^.Although no significant changes in the expression of OGT were observed in OGT^C921Y^ cortex (*p* = 0.788, *t*-test) (**Fig. S13A, D**), there was a significant increase in OGT/OGA protein ratio in OGT^C921Y^ mice (*p* = <0.0001, *t*-test) (**Fig. S13E**), which is driven by the reduction in OGA levels to compensate for reduced OGT activity. Despite this compensatory mechanism, global O-GlcNAcylation of proteins in the brain was drastically impaired in OGT^C921Y^ brain compared to WT (*p* = 0.0018, *t*-test) (**Fig. S13O, Q**). The perturbed regulation of *Ogt/Oga* ratio (Oga mRNA *p* = 0.0087; Ogt mRNA *p* = 0.0894; Ogt/Oga ratio *p* = <0.0001, *t*-test) was further confirmed at the transcriptional level by RT-PCR (**Fig. S14A, B, C**). Of note, similar O-GlcNAc dyshomeostasis at protein/mRNA levels was observed in other brain regions analysed except for OGT protein levels that were found reduced in both hippocampus and cerebellum (**Fig. S14**), suggesting brain region specific effect of the OGT^C921Y^ mutation. In the prefrontal cortex region, gene ontology analyses in the STRING database pointed to significant upregulation in the pathways regulating cellular processes (GO:0009987), cellular metabolic pathways (GO:0044237), nervous system development (GO:0007399), protein catabolic process (GO:0030163) and lysosomal transport (GO:0007041) (**Fig. 7C**). Among the top up regulated proteins identified in our dataset (**Fig. 7A**), thirty-five proteins belong to neurodevelopmental pathways, including proteins involved in neuronal migration (PLXND1, FAT3, ASTN2, NEUROD1), neurogenesis (NEUROD1, RBBP5, RBBP6) and synaptic function (CLCN3, SORCS3, AP3B1).

**Figure 7:**
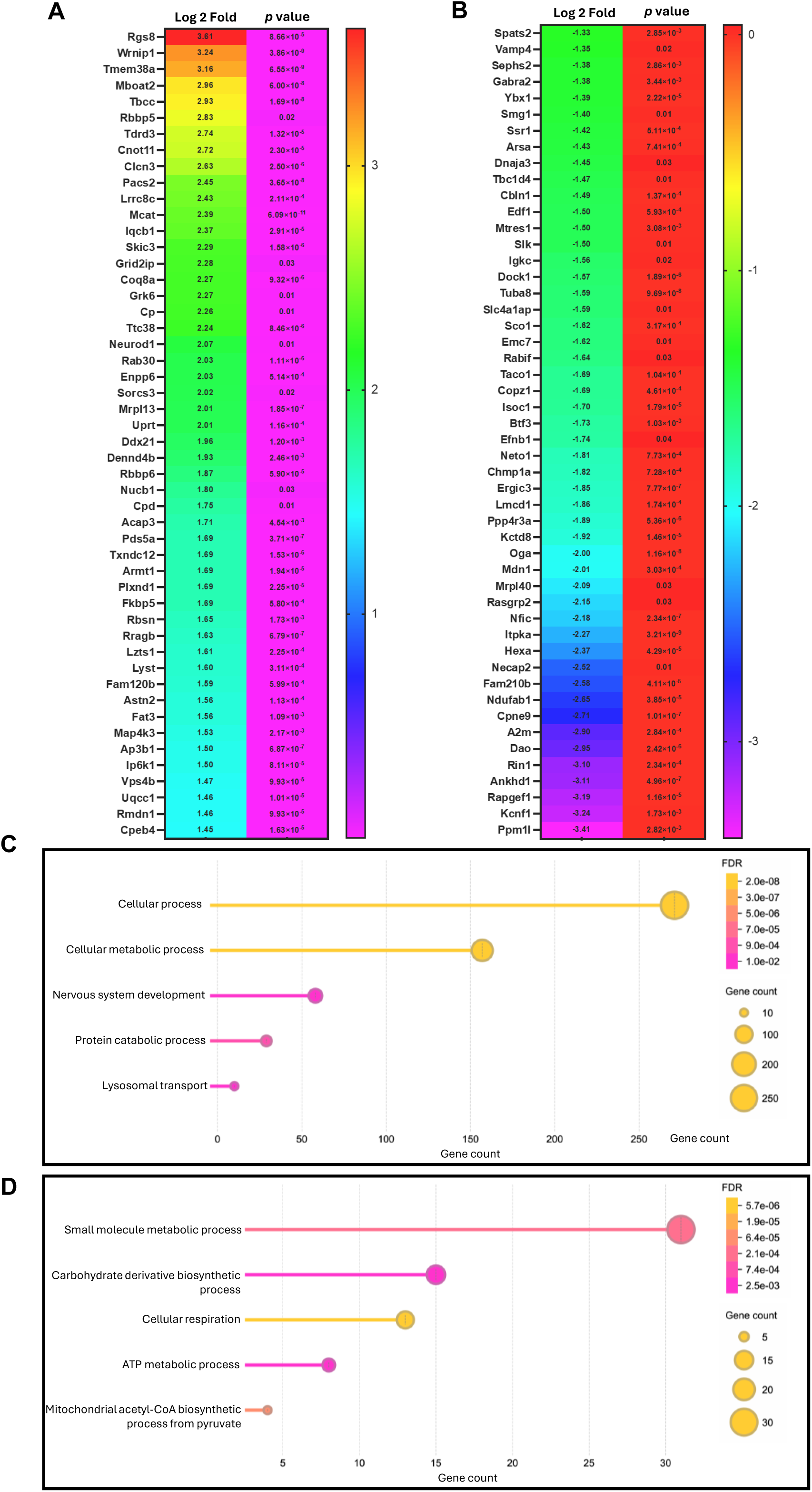
Quantitative proteomics analyses revealed distinctly perturbed molecular pathways in prefrontal cortex of the OGT^C921Y^ mice. (A) List of Top 50 proteins up-regulated (cut off criteria *p* value 0.05, log 2 change 1.45). (B) List of Top 50 proteins down-regulated (cut off criteria *p* value 0.05, log 2 change −1.33). (C) Up-regulated biological pathways enrichment in the STRING database. (D) Down-regulated biological pathways enrichment in the STRING database.

Intriguingly, we also identified significant downregulation in the pathways regulating small molecule metabolic process (GO: 0044281), carbohydrate derivative biosynthetic process (GO: 1901137), cellular respiration (GO: 0045333), ATP metabolic process (GO: 00046034) and mitochondrial acetyl-CoA biosynthetic process from pyruvate (GO:0061732) (**Fig. 7D**). Moreover, the perturbed proteomic signatures in the PFC corresponded to distinct Monarch enrichment profiles^86^ in the STRING database including abnormal CNS myelination (HP:0011400), agenesis of corpus callosum (HP: 0001274), upper motor neuron dysfunction (HP: 0002493) and abnormal muscle physiology and function (HP: 0011804; HP: 0003808; HP: 0001252; HP: 0001319; HP 0003394) (**Fig. S13A, B**).

Taken together, these data suggest that perturbations in protein O-GlcNAcylation in brain of OGT^C921Y^ are associated with distinct alterations in molecular pathways that potentially impact brain development.

## Discussion

Although several OGT variants linked to ID have been reported, the mechanisms underlying the disorder remain unknown. Potential mechanisms that have been proposed^63^ (**Fig. 8A**) include loss of O-GlcNAcylation on OGT substrates important for brain function and development^28,37,63^, OGT aggregation due to reduced OGT stability^63^, impaired OGT interactome^87^, misprocessing of HCF1^15^, or loss of OGA^27,28,37^, a common feature observed in models of OGT-ID. Lastly, O-GlcNAc dyshomeostasis has been recently suggested as a common mechanism in OGT-ID variants and has been proposed as a biomarker to identify new pathogenic variants using a stem cell reporter line^88^. Furthermore, these hypotheses are not mutually exclusive and can individually or in combination account for the cognitive dysfunction due to impaired brain development, defects in synaptogenesis/synaptic pruning and/or neural transmission impairment (**Fig. 8**). Vertebrate models are a key step towards deciphering the mechanisms linked to the disease, and here we have described a detailed characterization of a mouse model of one OGT-ID variant (OGT^C921Y^).

**Figure 8:**
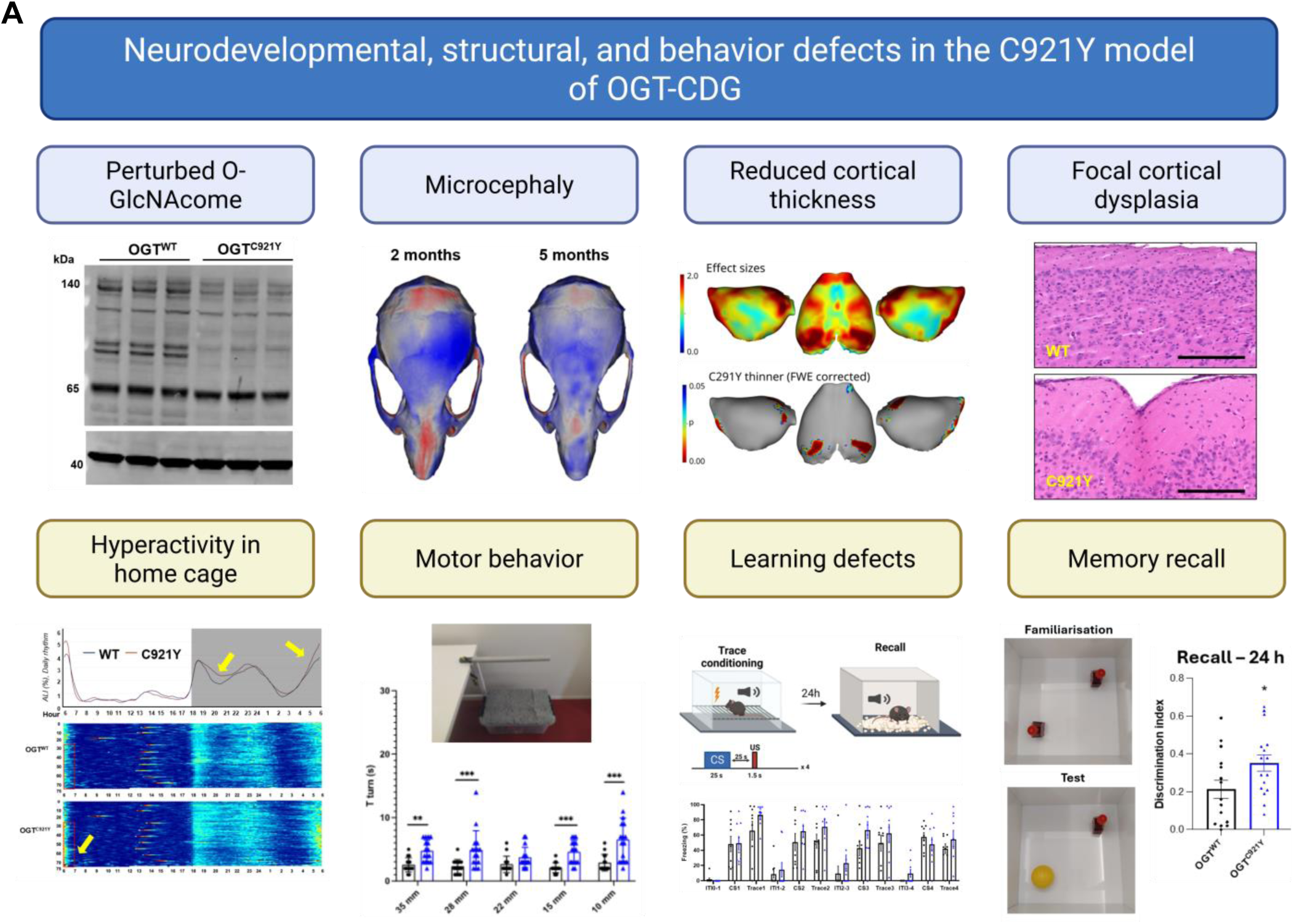
Graphical summary of the neurodevelopmental, structural and behaviour defects identified in the OGT^C921Y^ mice. This Figure was created using BioRender.com Sevillano quispe, O. (2026) https://BioRender.com/n21l133

The OGT^C921Y^ variant is found in a Danish family with three affected male siblings that inherited the variant through their affected mother. The male siblings exhibit dysmorphic features, ID with poor language skills and late onset epilepsy. Autistic features including repetitive mannerisms, and hyperactivity have also been reported^32^. Mild hyperactivity has also been reported in a patient carrying another OGT-ID variant^89^. The OGT^C921Y^ mice recapitulate features of impaired cognitive function, impulsive behaviour and increased spontaneous activity (or hyperactivity) both in their home cage and in the open field arena. These phenotypes (hyperactivity and impulsivity) are often observed in mouse models recapitulating human ID and autism disorders. For instance, *Fmr1^Ko^* mice exhibit hyperactivity in open field assays and increased exploratory behaviour as seen by an increased number of light-dark transitions in the dark-light paradigm similarly to our observations in the OGT^C921Y^ mice^90–92^. Moreover, hyperactivity and impulsivity behaviours present in a haploinsufficiency mouse model of *Kdm6b* were rescued with methylphenidate, a neurostimulant used to alleviate attention deficit and hyperactivity in individuals with ADHD^93^. In addition, the stereotypy and increased rearing activity observed in the OGT^C921Y^ mice suggest locomotor compulsive behaviour similar to that reported in the individuals carrying the C921Y variant^32^. We have observed that, unlike the WT mice, the locomotor activity of the OGT^C921Y^ mice remained unaffected during repeated measurements in the open field test, which could suggest an impairment in spatial habituation learning. Interestingly, defects in habituation learning were also observed in *Drosophila* models carrying OGT-ID variants during a light-off jump habituation test^94^. As habituation corresponds to the simplest form of learning^95^, these data hint towards some learning and memory defects in OGT^C921Y^ mice. This is further indicated by a delay in freezing behaviour during acquisition compared to WT littermates during the aversive Pavlovian paradigm, a well-established hippocampus- and cortex-dependent learning and memory.

Reduction of brain size has been reported in OGT-ID patients, including microcephaly and brain atrophy^29^. In the present study, both the shape and morphology of the skulls of the OGT^C921Y^ mice were affected. The morphological alteration of the skulls reduced the endocast volume (as an estimate of brain volume) significantly in both 2 month old and 5 month old OGT^C921Y^ mice compared to their WT littermates. MRI brain structural analyses indicate that redistribution of regional volumes rather than overall reduction underpins the lower total brain volume in the OGT^C921Y^ mice. The frontal lobes of the cerebral cortex were prominently affected and associated with bilateral reduced cortical thickness suggesting overall reduction of cortical grey matter. Reduced brain volume, in particular the prefrontal cortex region, has also been associated with several neurodevelopment disorders in human including ADHD^96,97^ and associated rodent models^98^. Moreover, reduced brain volume and grey matter density in cerebellum and cortical regions are observed in adolescents with ASD associated with intellectual deficits^96^.

While the MRI data suggest an overall reduction in the size of the prefrontal cortex, histological analyses revealed FCD and pseudosulci formation in areas of the cingulate (retrosplenial) cortex in the OGT^C921Y^ mice with patchy distribution of ectopic white matter in the vicinity of these cortical malformations. These features are reminiscent of cortical polymicrogyria seen in some types of CDG disorders such as muscular dystrophy-dystroglycanopathy (MDDG) syndromes^84,85^, and in some inborn errors of metabolism characterized by defective neural migration^99^. Given that OGT^C921Y^ mice exhibit considerable reduction in cortical protein O-GlcNAcylation, it is plausible to postulate that these forms of cortical dysplasia (psuedosulci) arise as neurodevelopmental sequelae to the perturbed O-GlcNAcome. FCD has been linked to cognitive impairment in certain X-linked intellectual disabilities (XLIDs), such as Fragile X syndrome^100^. FCD arises from localized malformations in the cortex caused by disruptions in neuronal proliferation, differentiation, and migration during corticogenesis^101–103^. However, some studies suggest that similar disruptions in the organization of cortical layers may also arise due to disorders of post-migrational development of neural progenitors and not solely due to defects in proliferation or migration^104^. Therefore, it remains to be determined whether the features of cortical malformation (microcephaly and disturbed cytoarchitecture in cingulate) in the OGT^C921Y^ mice pertain to these categories. Behavioural and neuroimaging studies indicate that distinct regions within the cingulate participate in complex cognitive tasks involving affective response and decision making^105,106^. Meta-analyses of neuroimaging studies in human subjects with psychiatric disease (major depressive disorder, bipolar disorder, schizophrenia, anxiety and addiction) purport grey matter loss in the dorsal anterior cingulate as a frequent finding^107,108^. Taken together, these suggest a link between the cortical structure defects and the behavioural deficits observed in the OGT^C921Y^ mice. However, further systemic interventions (e.g. optogenetic manipulations) will be needed to investigate whether the behavioural phenotypes of OGT^C921Y^ mice solely arise from cortical dysplasia in the cingulate cortex.

There is a significant dearth of information on possible cortical malformations in patients with OGT-ID. This is largely due to the lack of clinical data available as brain imaging has only been reported in a few cases, or that the defects are beyond the limits of detection/resolution in routine clinical MRI scans. Nevertheless, defects in white matter composition have been reported in patients with OGT-ID including thin corpus callosum and periventricular leukomalacia^29^. Both defects have previously been associated with learning difficulties and ID^96,109^. Changes in white matter structures have also been found in neuroimaging analysis of neurodevelopment disorders such as Fragile-X^110^, developmental delay^111,112^ and ADHD^113^. Together, these observations suggest that changes in white matter structures could underline learning and behavioural deficits in neurodevelopment disorders. Interestingly, the OGT^C921Y^ mice also showed reduced volumes of several white matter structures including corpus callosum, internal capsule, cerebral peduncle, corticospinal tracts and stria terminalis. These observations could guide future studies in investigating whether OGT-ID variants cause impaired myelination processes leading to cognitive impairment.

Our proteomics analyses revealed a number of perturbed molecular pathways in the prefrontal cortex of the OGT^C921Y^ mice. Interestingly, some of the deregulated proteins are associated with brain malformation including progressive microcephaly (ARSA, CHMP1A, QARS1)^114–116^, cortical dysplasia (RALGAPB, PACS2)^90,9^, myelination defects (ARSA, CHMP1A, GABRA2, HEXA, RMND1)^114,117–121^ and agenesis of the corpus callosum (EFNB1, TUBA8, CLCN3, PLXND1)^122–125^, providing potential candidates underlying the brain defects identified in the OGT^C921Y^ mice. In addition, the RHO GTPase cycle pathway (HAS-9012999) was significantly upregulated from the Reactome database. RHO GTPase family are involved in cell migration, division and polarity and play crucial roles in neurodevelopment^126^. While a detailed description of all the top 50 upregulated proteins is beyond the scope of this Discussion, the ubiquitin ligase RB binding protein 6 (RBBP6) promotes ubiquitination of the Y box protein 1 (YBX1) followed by its degradation by the proteosome. YBX1 is a nucleic acid binding protein required for forebrain specification, cell proliferation and neuronal differentiation through the suppression of RNA polymerase II mediated transcription^127^. Interestingly, while RBBP6 was found up regulated, its target YBX1 was identified as significantly down regulated in our dataset, suggesting a role of the RBB6P/YBX1 axis in the brain structural defects observed in the OGT^C921Y^ mice. Future studies at early timepoints during brain development will be needed to explore these candidate conveyors of OGT-ID.

Whereas structural brain defects suggest a neurodevelopment origin, it is as yet unknown whether the associated cortical dysfunction originates from structural defects during neurodevelopment or neurophysiological defects due to the continuous presence of the catalytically impaired OGT^C921Y^ variant in neurons, their synapses and glia. Non-invasive monitoring of the OGT^C921Y^ mice activity patterns (in DVC) showed that spontaneous hyperactivity was not detected from weaning but continuously progressed from approximatively 7 weeks of age, coinciding with a delay in postnatal growth. With reasonable caution, this is reminiscent of clinical findings in patients with OGT-ID who generally start developing symptoms in the first two years. From a translational perspective, these features hint towards a progressive postnatal appearance of behaviour and morphological deficits in the OGT^C921Y^ mice, thus potentially offering a therapeutic window for future interventions. It is hoped that cognitive impairment due to prefrontal cortical dysfunction can partly be rescued postnatally even in presence of brain structural defects, with notable examples from research in models of Fragile X and Retts neurodevelopmental syndromes ^69,97,100^.

## Conclusions

In conclusion, we report that O-GlcNAc dyshomeostasis in brains of OGT^C921Y^ mice is associated with distinct behavioural phenotypes reflecting hyperactivity, impulsivity and learning deficits. These phenotypes were accompanied by features consistent with perturbed neurodevelopment including skull deformation, microcephaly and focal cortical dysplasia in the cingulate cortex (**Fig. 8B**). Moreover, the glimpse offered by changes in the neocortical proteome of OGT^C921Y^ mice and the observed O-GlcNAc dyshomeostasis will guide future studies in unravelling the pathophysiology of the disorder, and hold promise for the development of novel therapeutic interventions in OGT-ID.

## Data availability

Data will be available upon request.

## Acknowledgements

The authors would like to thank Trine Mikkelesen (JRN lab) for the assistance with the histology workflow, Kristian Graff (Department of Molecular Biology and Genetics, Aarhus University), for the assistance in behaviour equipment build-up and Kamilla Zahll Hornbek (Department of Biomedicine, Aarhus University) for breeding and animal care assistance.

## Author contributions

F.A. and D.M.F.v.A conceived the study; F.A., I.F., C.S.S., S.T.B., K.S.C., B.A., J.S.T. performed experiments; C.S. performed mass spectrometry; F.A., I.F., C.S.S., K.S.C., I.E.A, S.F.E., B.H. analysed data; A.J. and J.R.N. performed histology and image analyses; O.G.S.Q. provided illustration used in Figure 8; F.A., A.J. and D.M.F.v.A. interpreted the data and wrote the manuscript with input from all authors.

## Supplementary material

**Supplementary Figure S1:**
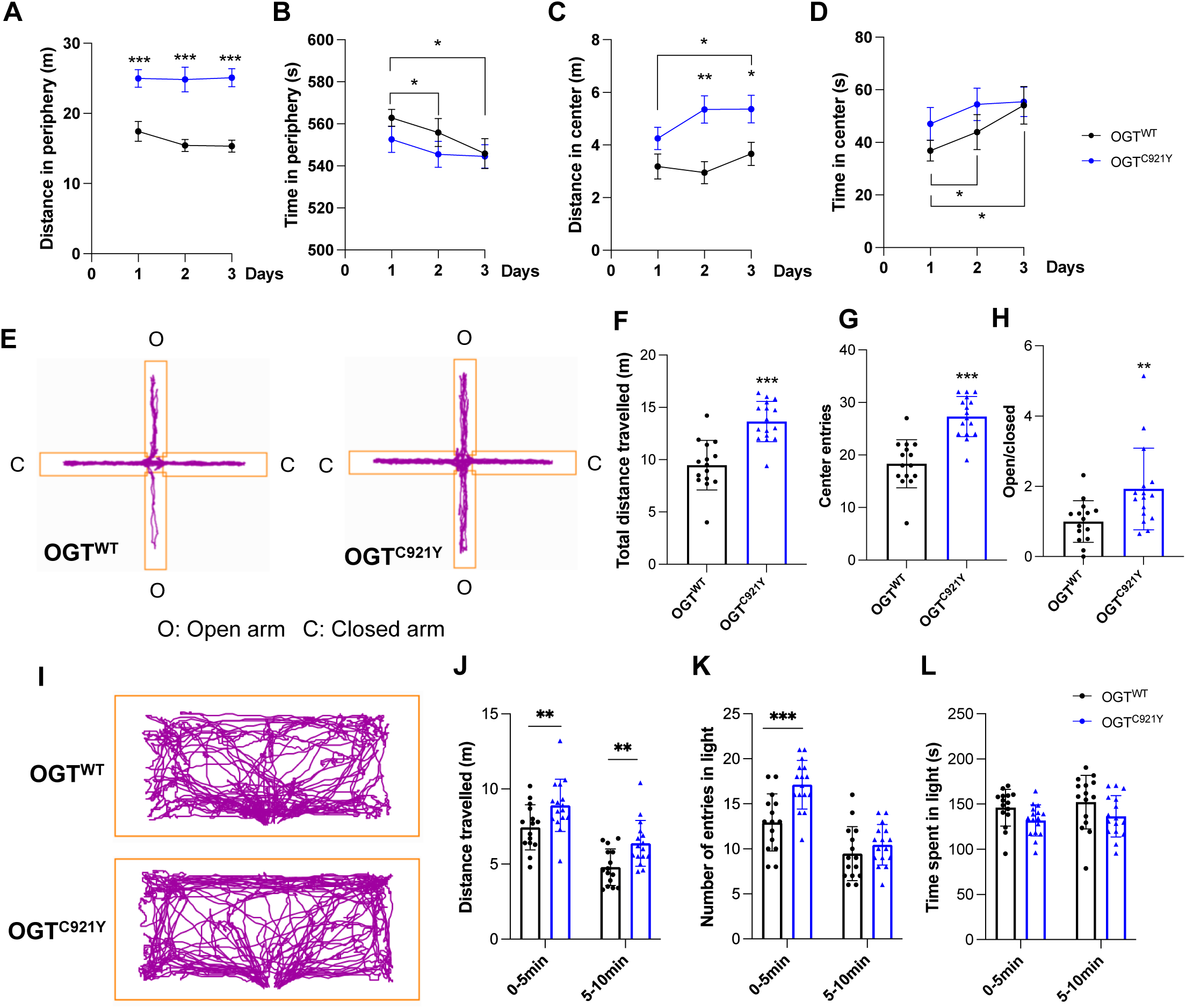
OGT^C921Y^ mice show similar anxiety-like behaviour and increased exploration. Significance is shown as \**p* < 0.05, \*\**p* < 0.01 and \*\*\**p* < 0.001. (A) Distance travelled in periphery by male OGT^WT^ (n = 15) and OGT^C921Y^ (n = 16) mice in the open field arena. Two-way ANOVA (Alpha, 0.05) followed by Tukey’s column comparisons. (B) Time spent in periphery by male OGT^WT^ (n = 15) and OGT^C921Y^ (n = 16) mice in the open field arena. Two-way ANOVA (Alpha, 0.05) followed by Tukey’s column comparisons. (C) Distance travelled in centre by male OGT^WT^ (n = 15) and OGT^C921Y^ (n = 16) mice in the open field arena. Two-way ANOVA (Alpha, 0.05) followed by Tukey’s column comparisons. (D) Time spent in centre by male OGT^WT^ (n = 15) and OGT^C921Y^ (n = 16) mice in the open field arena. Two-way ANOVA (Alpha, 0.05) followed by Tukey’s column comparisons. (E) Representative tracking plot of male OGT^WT^ and OGT^C921Y^ mice in the elevated plus maze (EPM). (F) Total distance travelled of male OGT^WT^ (n = 15) and OGT^C921Y^ (n = 16) mice in the EPM. Student *t* test was used for statistics. (G) Number of centre entries of male OGT^WT^ (n = 15) and OGT^C921Y^ (n = 16) mice in the EPM. Student *t* test was used for statistics. (H) Ratio between time spent in open and close arms by male OGT^WT^ (n = 15) and OGT^C921Y^ (n = 16) mice in the EPM. Student *t* test was used for statistics. (I) Representative tracking plot of male OGT^WT^ and OGT^C921Y^ mice in the light compartment during the dark-light paradigm (D/L). (J) Total distance travelled of male OGT^WT^ (n = 15) and OGT^C921Y^ (n = 16) mice in the light compartment during the dark-light paradigm (D/L). Two-way ANOVA (Alpha, 0.05) followed by Tukey’s column comparisons. (K) Number of centre entries of male OGT^WT^ (n = 15) and OGT^C921Y^ (n = 16) mice in the light compartment during the dark-light paradigm (D/L). Two-way ANOVA (Alpha, 0.05) followed by Tukey’s column comparisons. (L) Time spent of male OGT^WT^ (n = 15) and OGT^C921Y^ (n = 16) mice in the light compartment during the dark-light paradigm (D/L). Two-way ANOVA (Alpha, 0.05) followed by Tukey’s column comparisons.

**Supplementary Figure S2:**
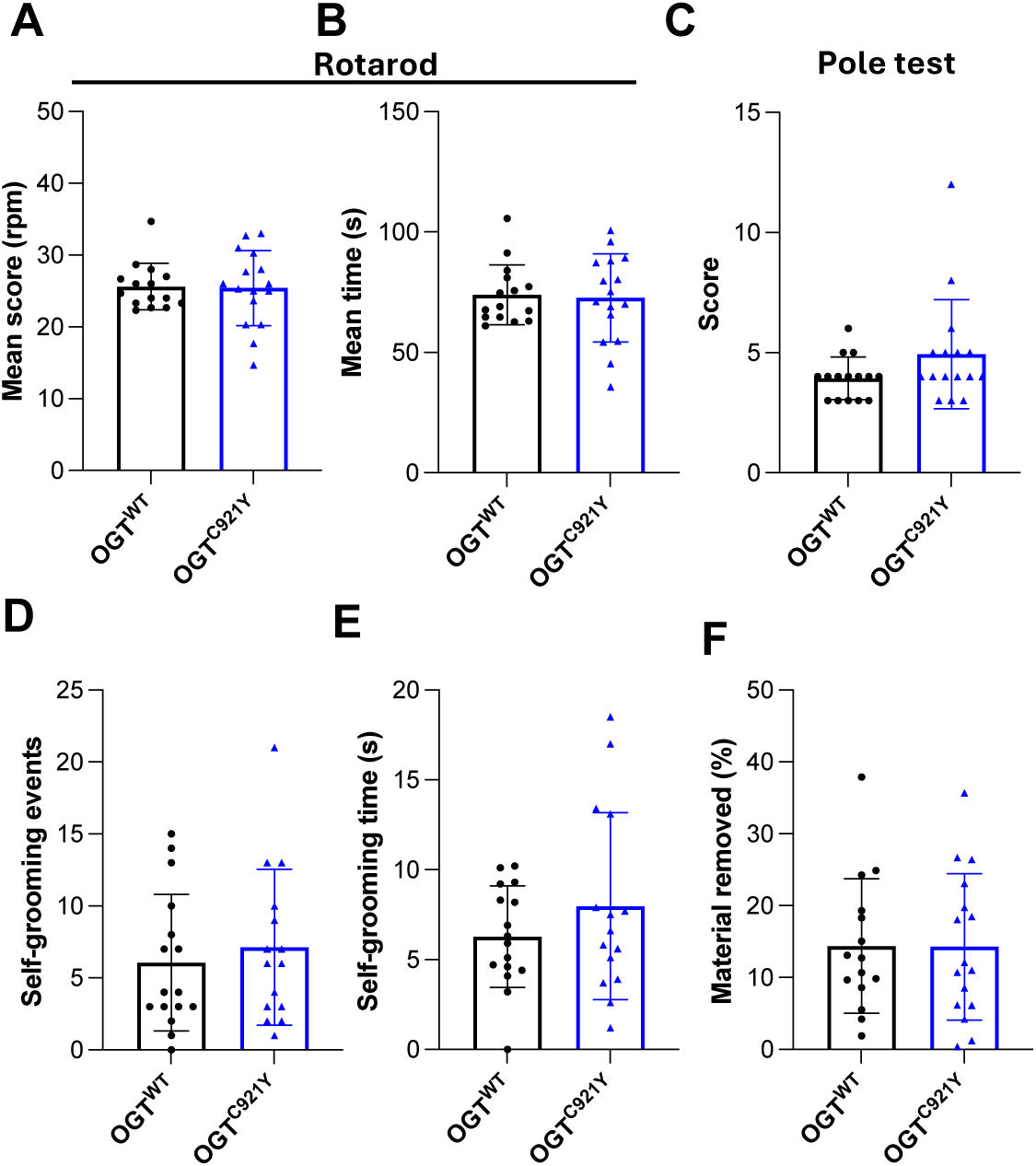
OGT^C921Y^ mice show similar motor skills. Significance is shown as \**p* < 0.05, \*\**p* < 0.01 and \*\*\**p* < 0.001. (A) Mean score reached by male OGT^WT^ (n = 15) and OGT^C921Y^ (n = 16) mice on the rotarod. (B) Mean time spent by male OGT^WT^ (n = 15) and OGT^C921Y^ (n = 16) mice on the rotarod. (C) Score reached by male OGT^WT^ (n = 15) and OGT^C921Y^ (n = 16) mice during the pole test. (D) Number of self-grooming events male OGT^WT^ (n = 15) and OGT^C921Y^ (n = 16) mice during 3 min observation. (E) Time spent self-grooming by male OGT^WT^ (n = 15) and OGT^C921Y^ (n = 16) mice during 3 min observation. (F) Percentage of material removed from a cotton pad by male OGT^WT^ (n = 15) and OGT^C921Y^ (n = 16) mice during 3 min during the nesting test.

**Supplementary Figure S3:**
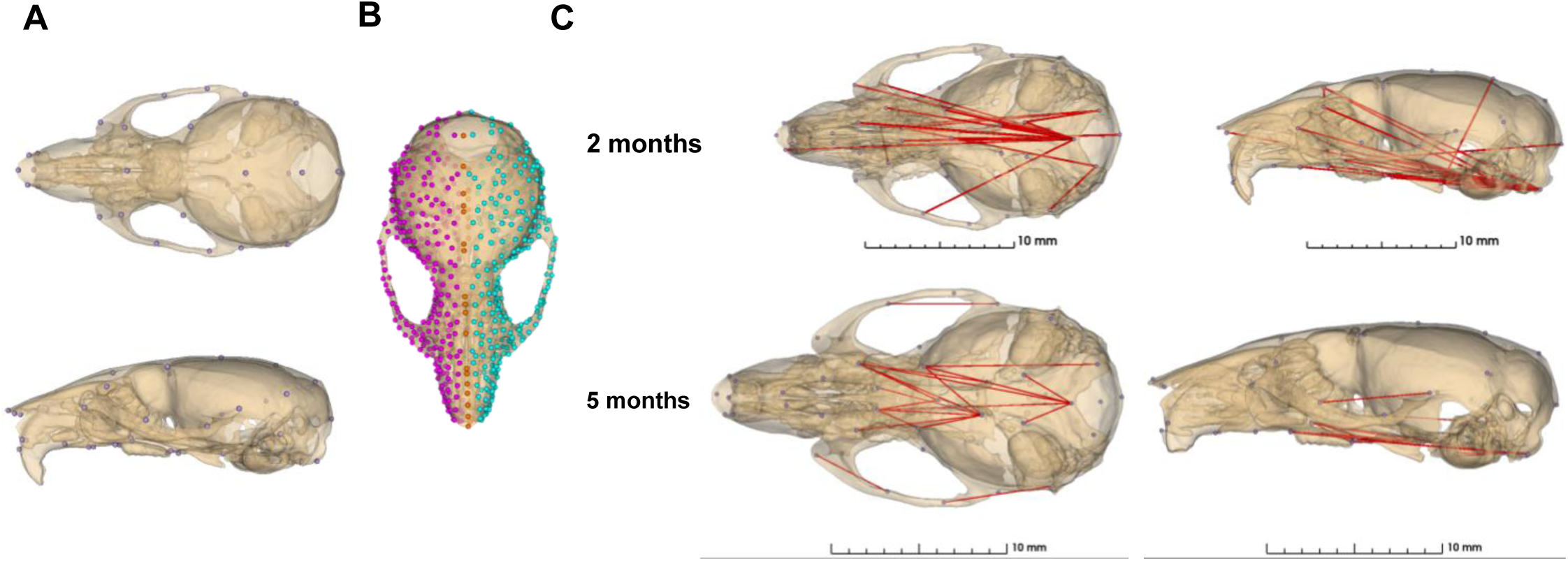
OGT^C921Y^ mice show reduced distances in the cranial base. (A) Lateral and superior views of a representative microCT 3D reconstruction of a mouse skull presenting the 45 landmarks used for Euclidean distance matrix analysis (EDMA) (left) and surface markups used for Principal Components Analysis and average shapes generation (right). (B) Lateral and inferior views of representative 3D reconstructions of mouse skulls presenting the linear distances that are at least 5% shorter in 2 and 5 months old OGT^C921Y^ skulls (p < 0.01, two-tailed unpaired t-test used)

**Supplementary Figure S4:**
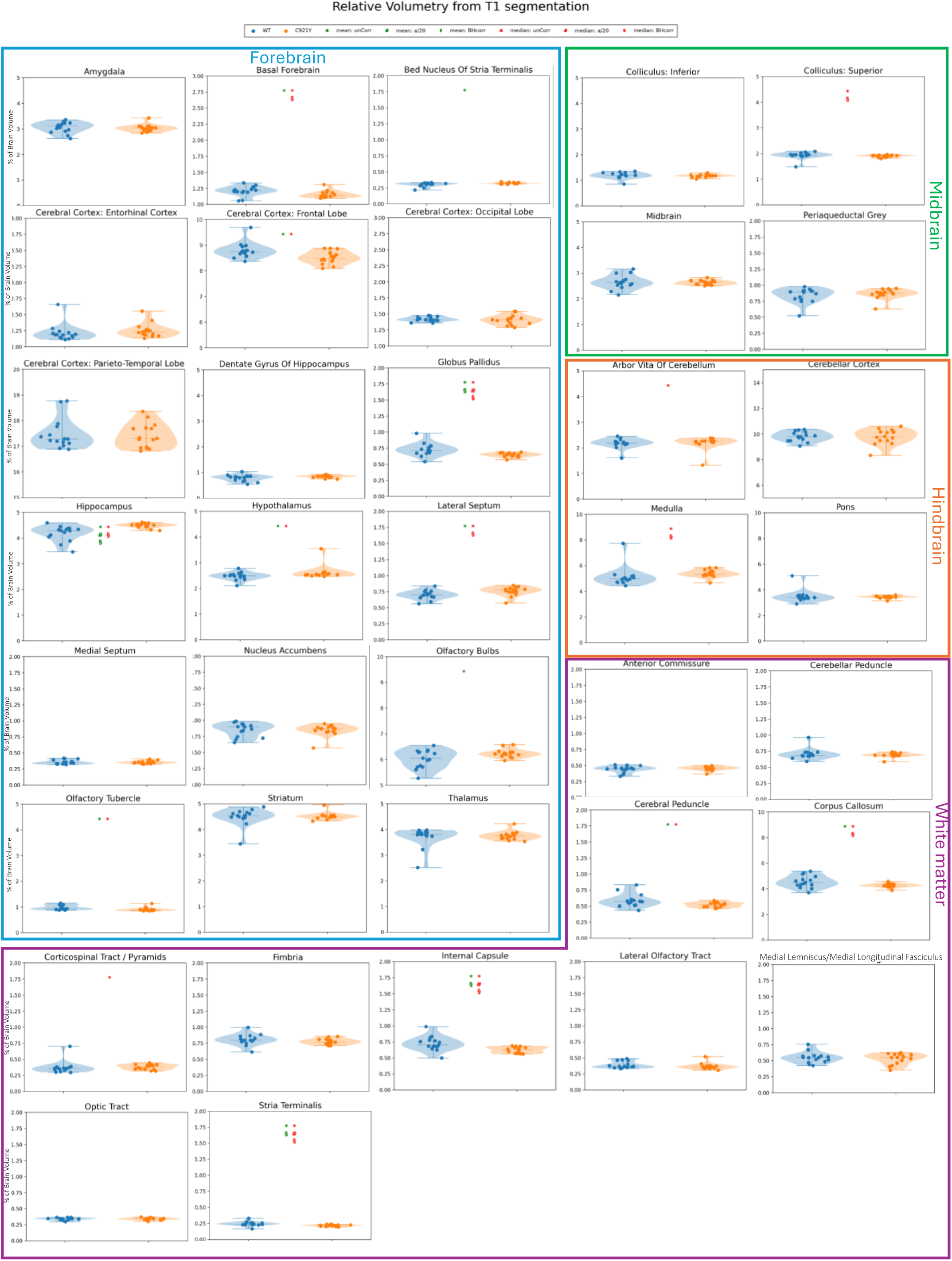
Volumetry from segmentation of T1-weighted structural images (FLASH) from Magnetic Resonance Imaging (MRI) illustrated as absolute volumes. Asterisks (*) indicate uncorrected significance (*p* < 0.05), based on permutation tests (100k permutations for each region). Pound symbols (#) indicate significance below alpha (0.05) divided by total number regions tested (40) for mean (green) and median (red), respectively. Section signs (§) indicate significance (*p* < 0.05) with *p* values adjusted for false discovery rate (Benjamini-Hochsberg, BH). Y-axes are scaled to individual ROIs to highlight group variation and difference.

**Supplementary Figure S5:**
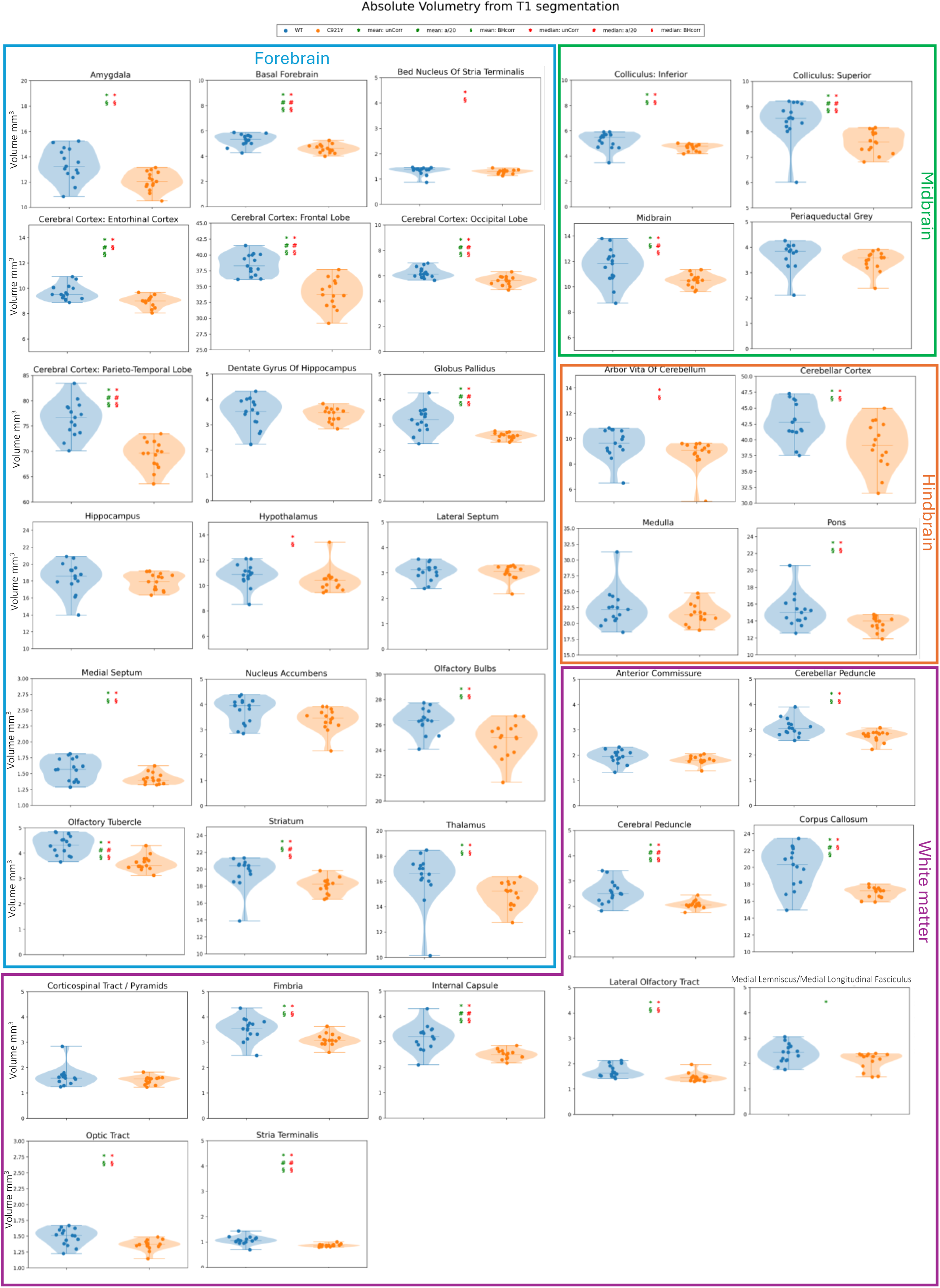
Relative volumetry from T1 segmentation. Volumetry from segmentation of T1-weighted structural images (FLASH) from Magnetic Resonance Imaging (MRI) illustrated as regional relative volumes (divided by individual total brain volume). Asterisks (*) indicate uncorrected significance (*p* < 0.05), based on permutation tests (100k permutations for each region). Pound symbols (#) indicate significance below alpha (0.05) divided by total number regions tested (40) for mean (green) and median (red), respectively. Section signs (§) indicate significance (*p* < 0.05) with *p* values adjusted for false discovery rate (Benjamini-Hochsberg, BH). Y-axes are scaled to individual ROIs to highlight group variation and difference.

**Supplementary Figure S6:**
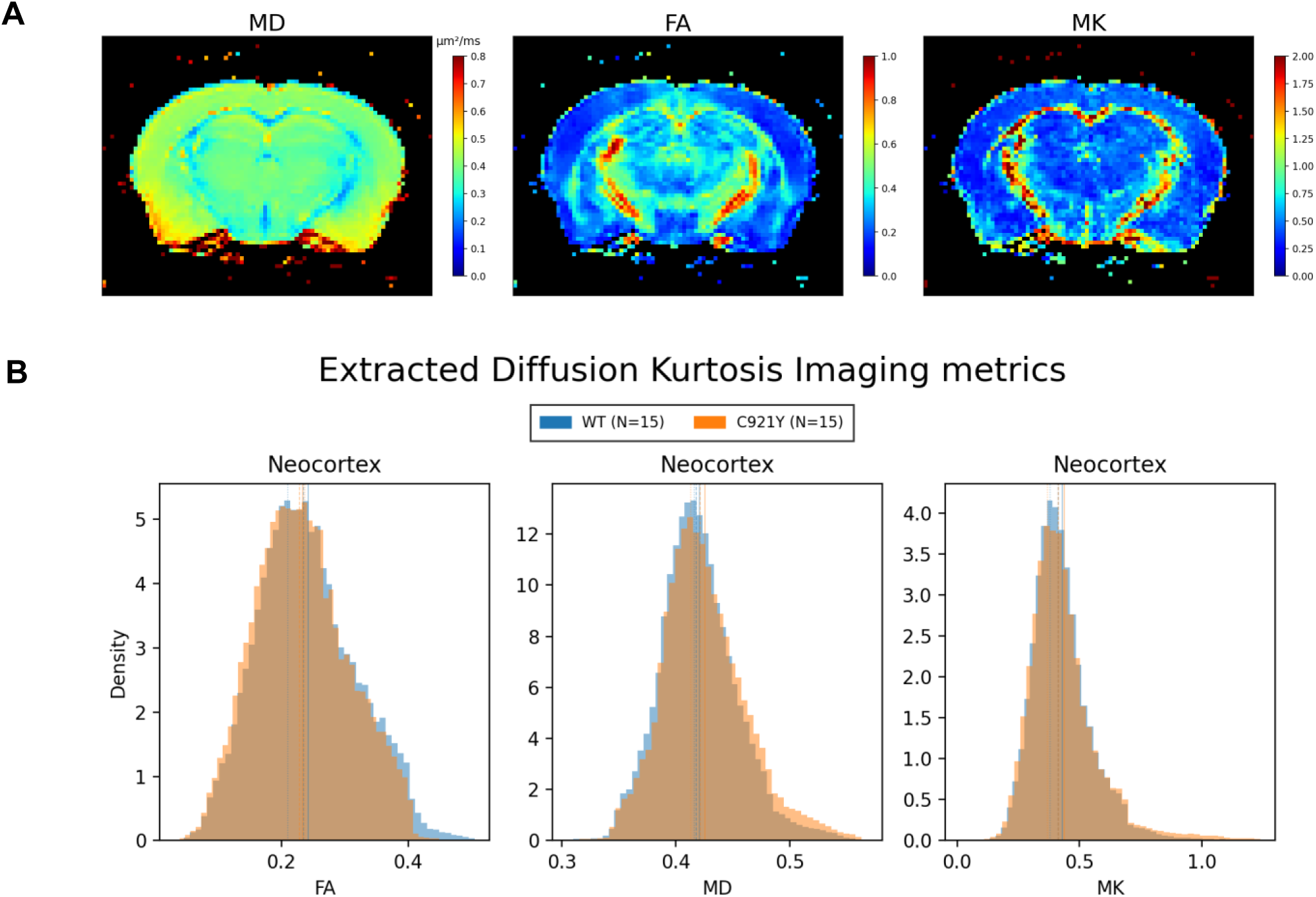
OGT^C921Y^ mice show conserved DKI metrics for Neocortex. (A) Middle coronal section of subject:207 illustrating the different calculated DKI metrics: Mean Diffusivity (MD), Fractional Anisotropy (FA) and Mean Kurtosis (MK). (B) Extracted DKI metrics (MD, FA and MK) for Neocortex of all subjects in each group for neocortex of all mice.

**Supplementary Figure S7:**
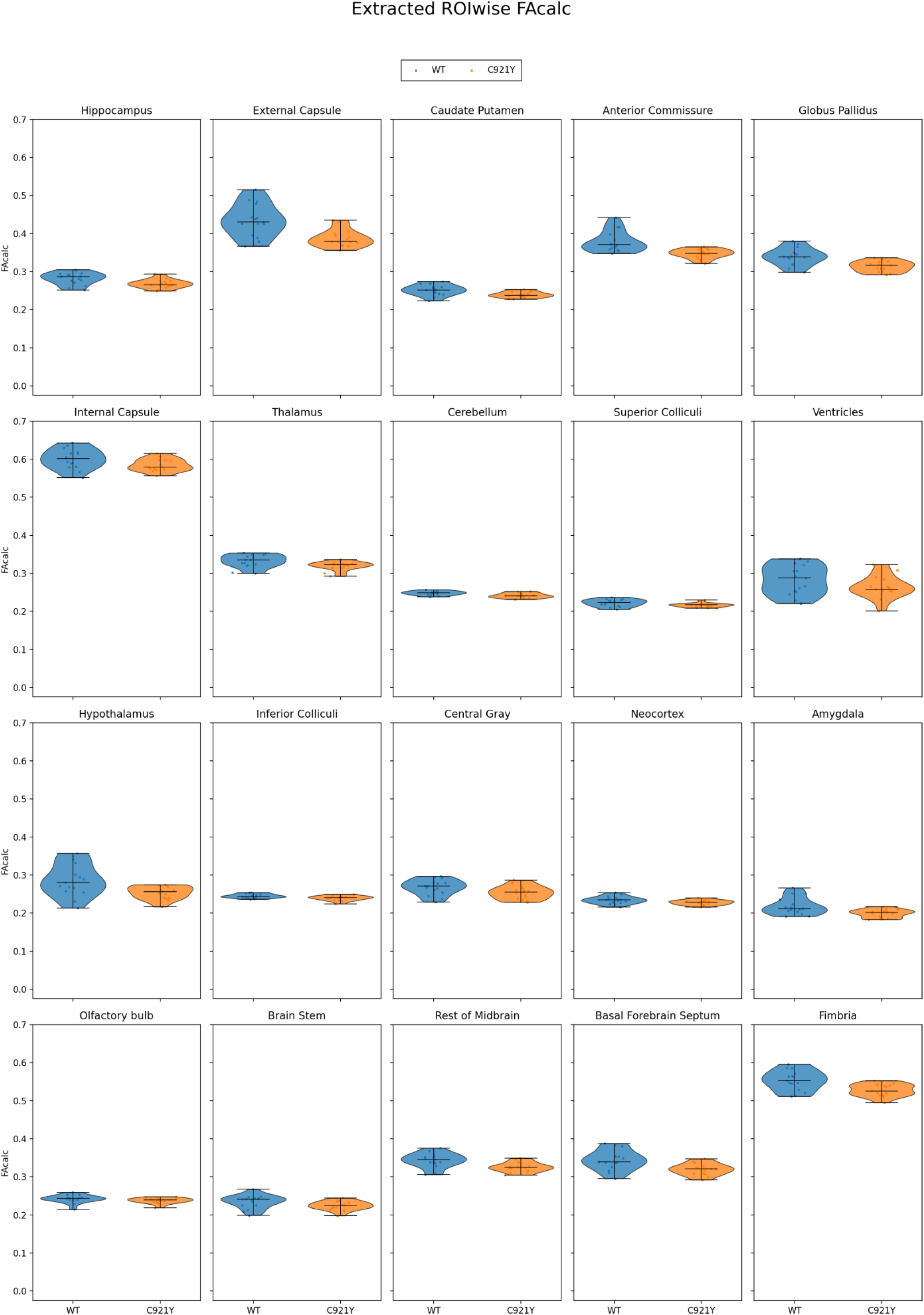
**Subject-wise median of extracted Fractional Anisotropy (FA) values in 20 selected regions of interest (ROI).**

**Supplementary Figure S8:**
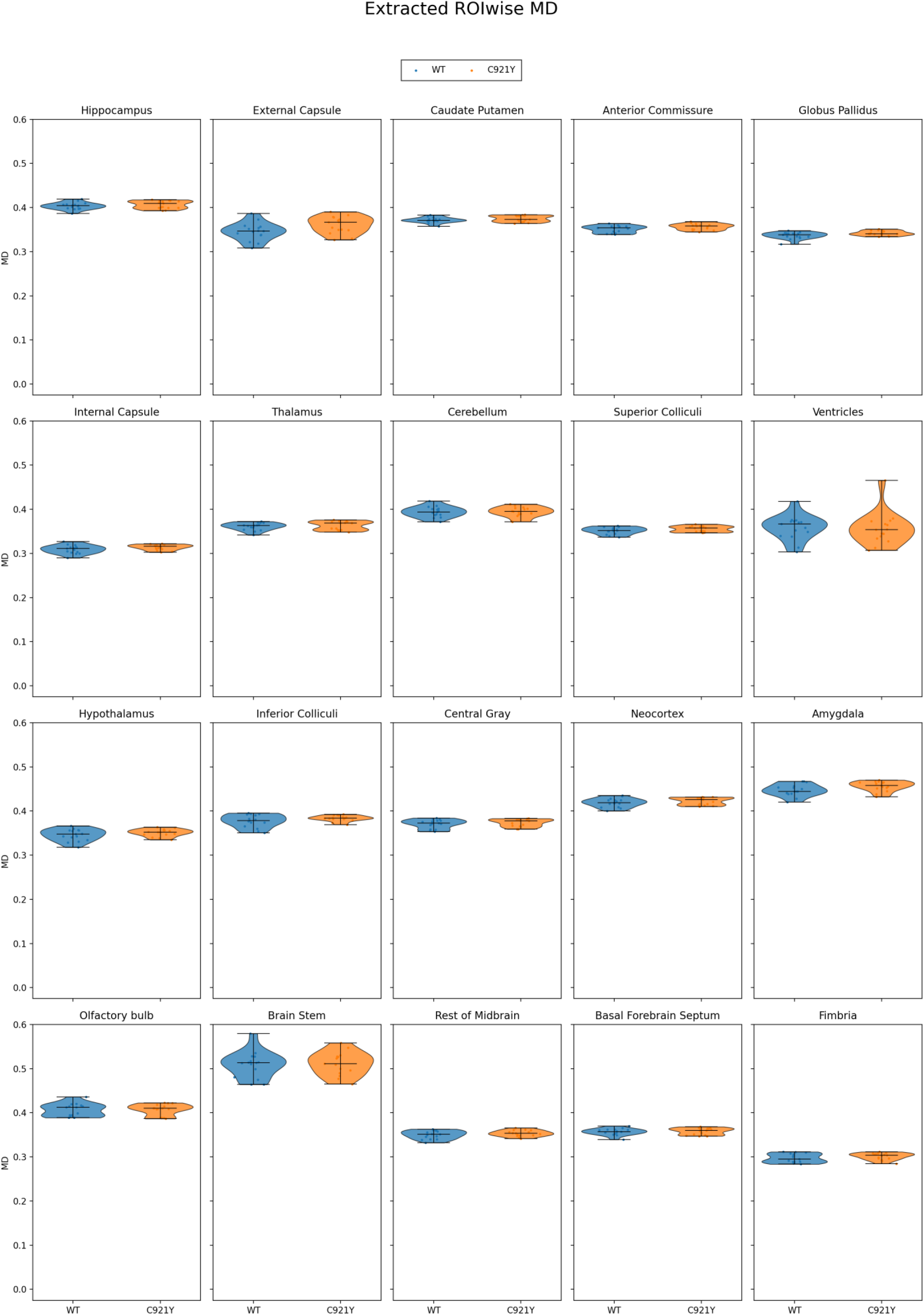
**Subject-wise median of extracted Mean Diffusivity (MD) values in 20 selected regions of interest (ROI).**

**Supplementary Figure S9:**
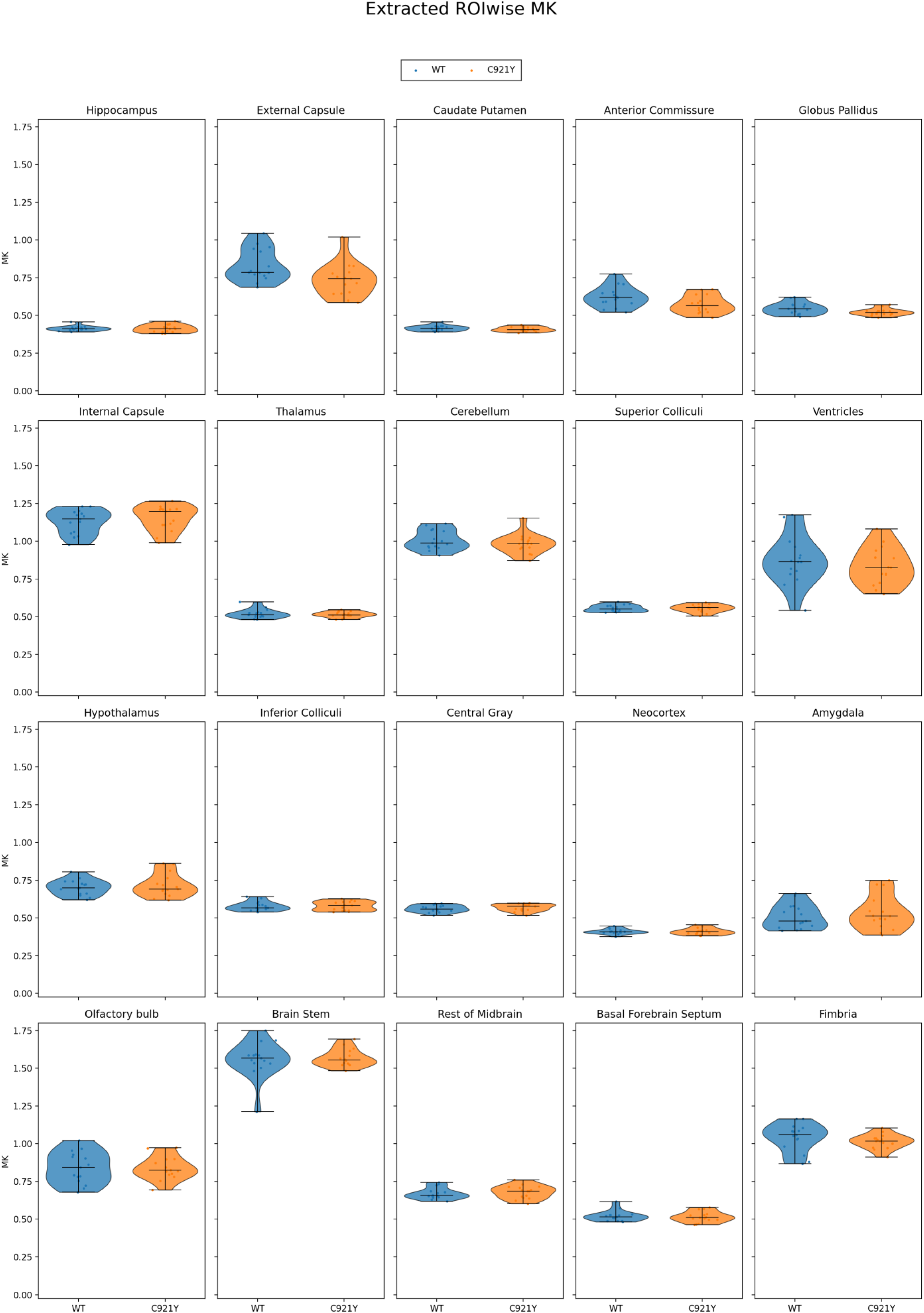
**Subject-wise median of extracted Mean Kurtosis (MK) values in 20 selected regions of interest (ROI).**

**Supplementary Figure S10:**
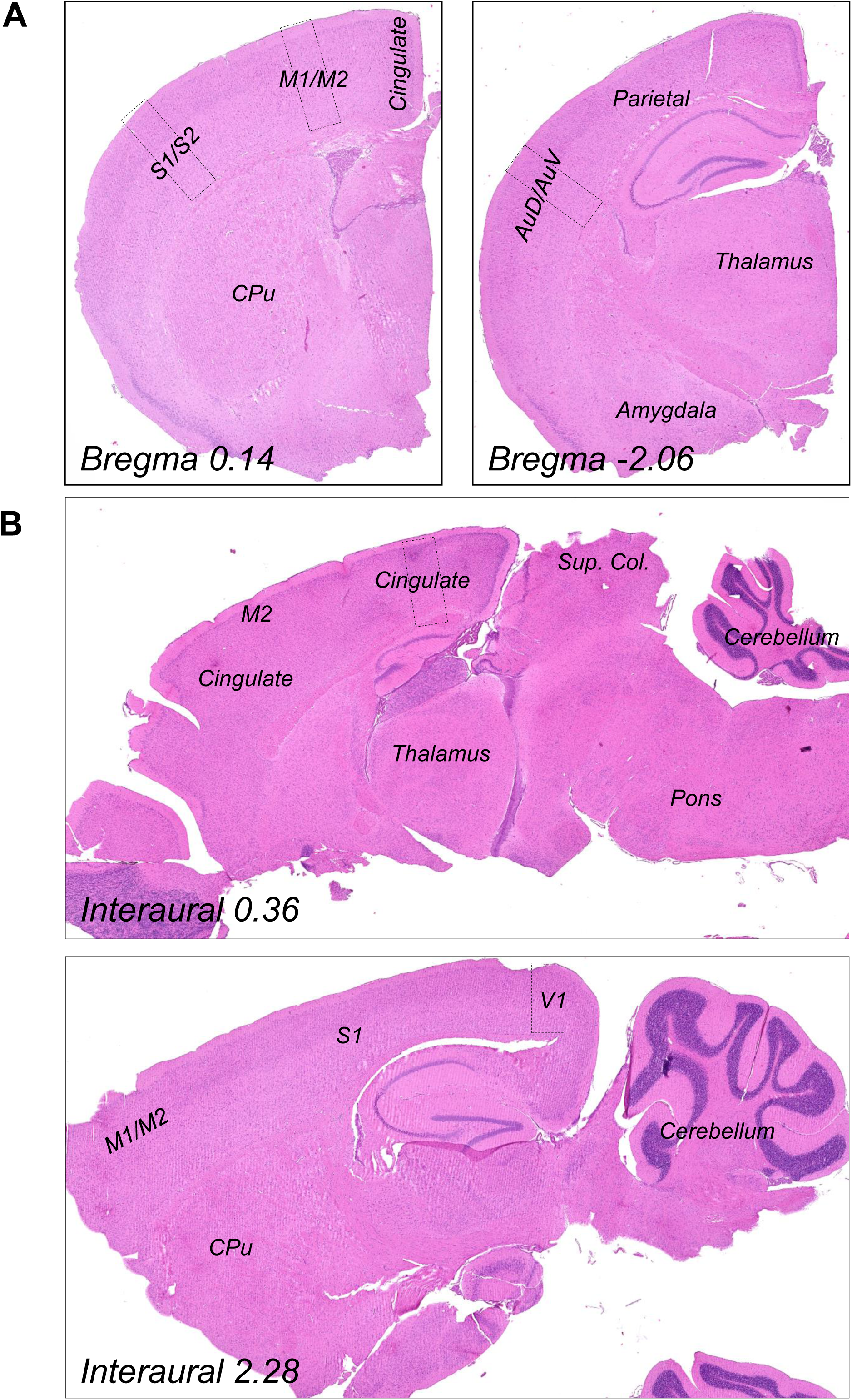
Panoramic views of the H&E stained sagittal and coronal brain sections depicting neuroanatomical landmarks used for measurements of cortical thickness and cell density. (A) Representative images showing coronal panoramic views of the brain sections used for the measurements of cortical cell density. The sections were used to sample regions in the cortex at the level of Begma 0.14 and Bregma −2.06. Regional cell counts were obtained from primary motor cortex M1/M2 and primary somatosensory cortex S1/S2, and from the auditory cortex. (B) Representative images showing sagittal panoramic views of the brain sections used for the measurements of cortical cell density. The sections were used to sample medial (interaural, 0.36, in C) and lateral (interaural, 2.28, in D) regions of the cortex. Regional cell counts were obtained from cingulate cortex and visual cortex V. Abbreviations: AuD/AuV, auditory cortex (in B); CPu, caudate putamen; superior colliculus. Neuroanatomical annotations are based on the Paxinos and Franklin’s Mouse Brain in Stereotaxic Coordinates, Elsevier Publishing, 4th Edition.

**Supplementary Figure S11:**
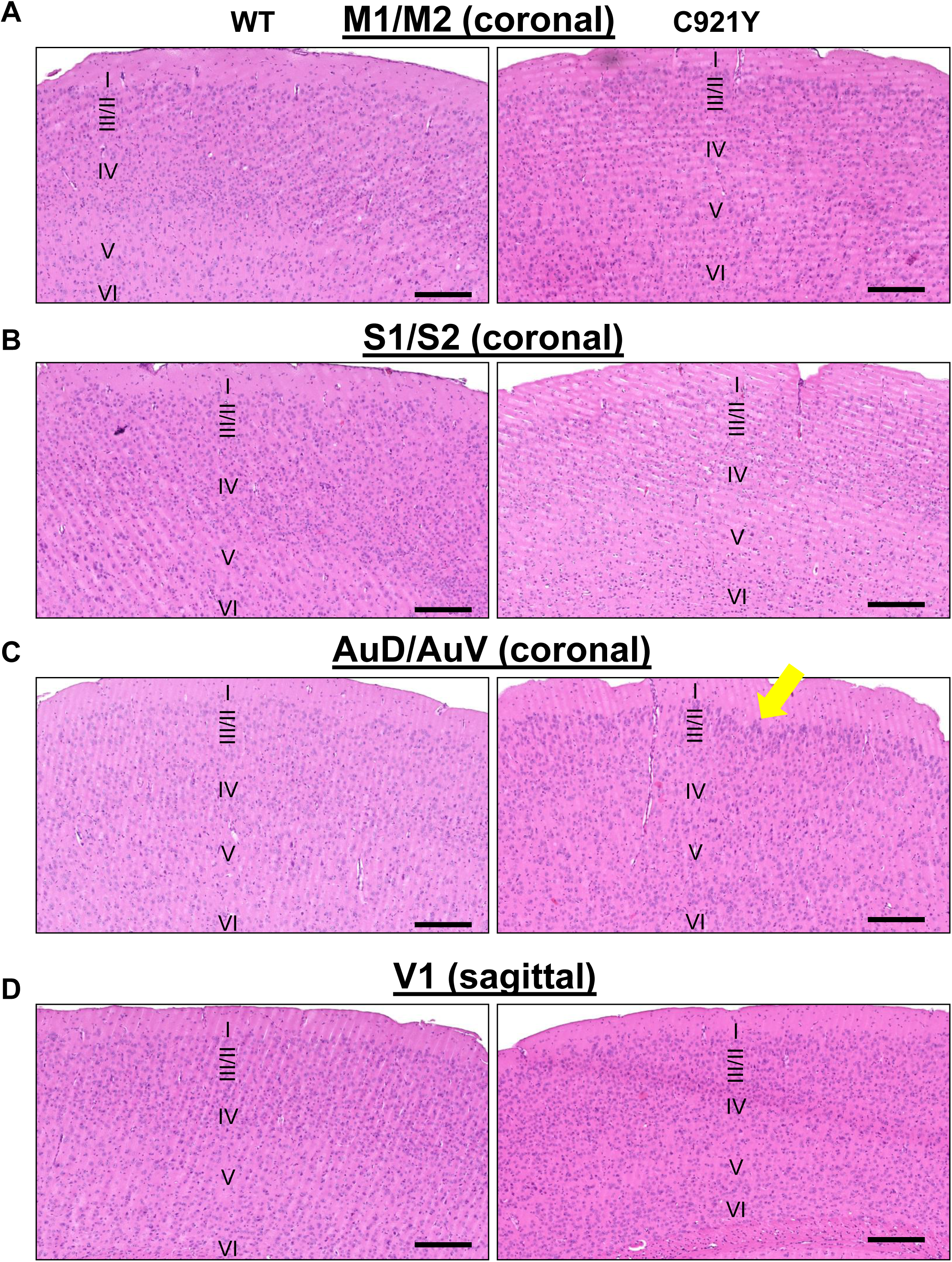
Low magnification (10X) views of the H&E stained sagittal and coronal brain sections used for assessing cortical organization. Representative images showing low magnification (10X) views of select cortical regions from OGT^WT^ (in A) and OGT^C921Y^ (in B) mice. Areas shown include primary motor cortex M1/M2, primary somatosensory cortex S1/S2, auditory cortex and visual cortex V1. Notice a mild degree of cortical dysplasia in layers II-III in the auditory cortex of OGT^C921Y^ mice (yellow arrow). The images were obtained from coronal sections (Bregma, 0.14, for M1/M2 and S1/S2; Bregma, −2.06, auditory cortex) and sagittal sections (Interaural 2.28, visual cortex V1). Scale bar = 200 µm. Also see Fig. S10. Neuroanatomical annotations are based on the Paxinos and Franklin’s Mouse Brain in Stereotaxic Coordinates, Elsevier Publishing, 4th Edition.

**Figure S12.**
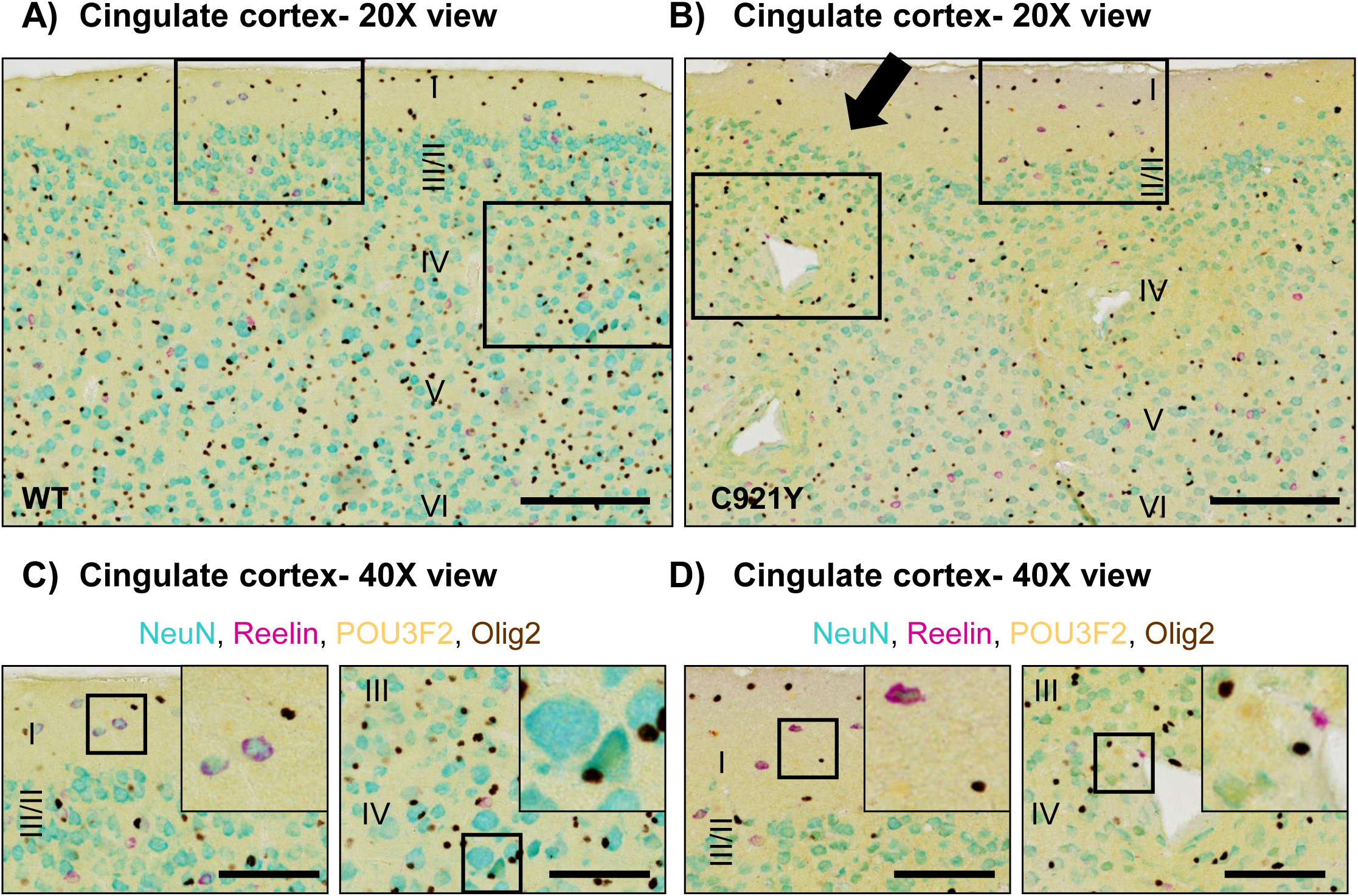
Histological characterization of cortical malformations by multiplex IHC. (A-B) Representative low magnification (20X) sagittal views of WT (A) and OGT^C921Y^ mice (B), showing cellular populations in the vicinity of nodular cortical malformation in the cingulate cortex of the latter using multiplex IHC chromogenic detection (NeuN in teal, Olig2 in brown, Reelin in purple and POU3F2 in yellow)- scale bar = 200 µm. Numerals I-VI in S12a-d indicate neocortical layers. (C-D) Representative high magnification (40X) views extracted from the insets in A (wild type) and B (OGT^C921Y^) respectively. Notice the undulating pattern of NeuN in layers II/III (black arrow, in B) and mixed cellular populations in the vicinity of cortical malformations in layers II-V of an OGT^C921Y^ animal-scale bar = 100 µm.

**Supplementary Figure S13:**
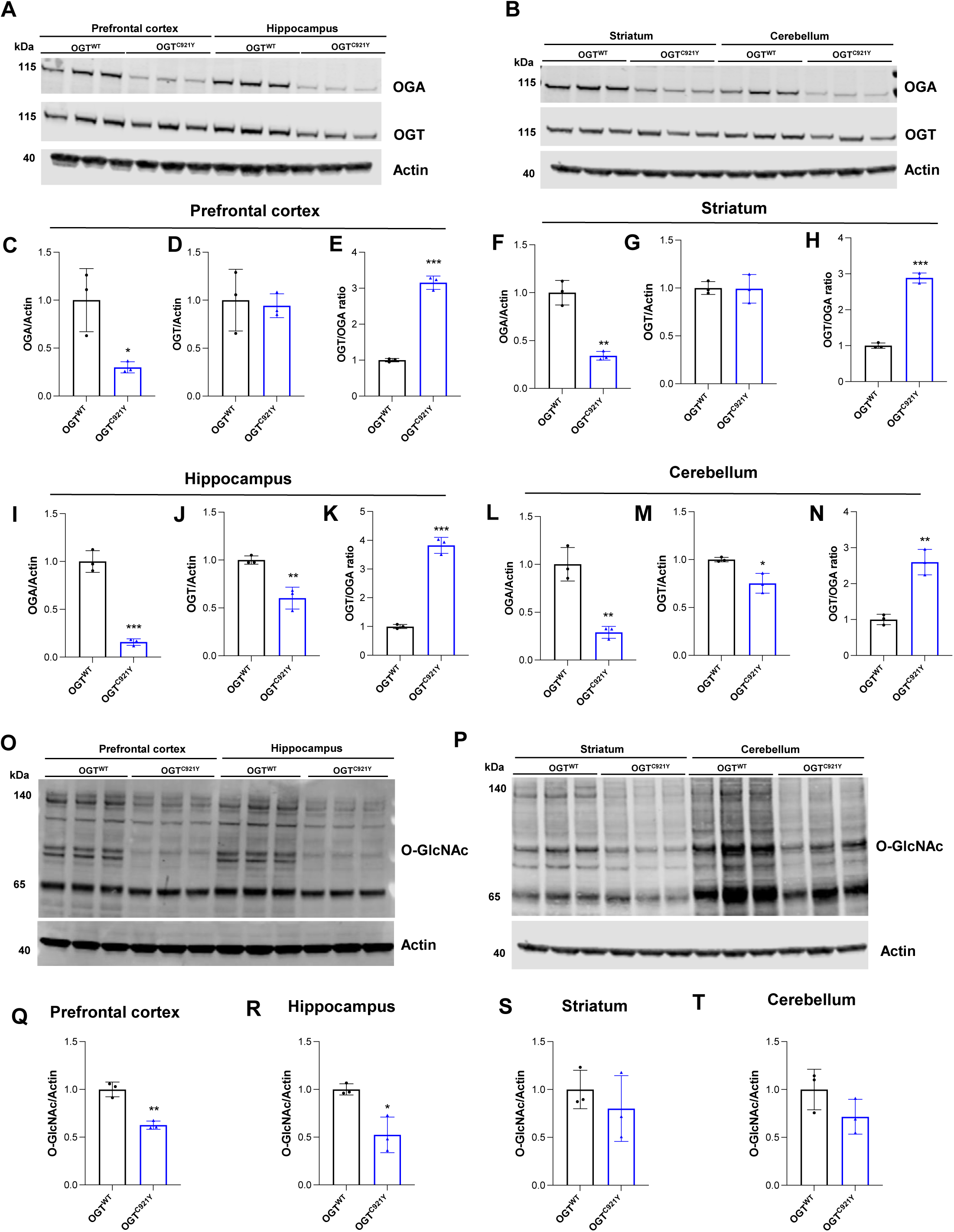
OGT^C921Y^ mice show altered OGT/OGA ratio and global reduction in O-GlcNAc levels in brain regions. Significance is shown as * *p* < 0.05, ** *p* < 0.01 and *** *p* < 0.001 (n = 3 per group). Student *t* test was used for statistics. (A) Western blot of OGA and OGT level in male prefrontal cortex and hippocampus of OGT^WT^ and OGT^C921Y^ mice. Actin antibodies were used as loading control. (B) Western blot of OGA and OGT level in male striatum and cerebellum of OGT^WT^ and OGT^C921Y^ mice. Actin antibodies were used as loading control. Quantification of OGA (C), OGT (D) and OGT/OGA ratio (E) proteins levels in prefrontal cortex from the Western blot in panel A. Quantification of OGA (F), OGT (G) and OGT/OGA ratio (H) proteins levels in striatum from the Western blot in panel B. Quantification of OGA (I), OGT (J) and OGT/OGA ratio (K) proteins levels in hippocampus from the Western blot in panel A. Quantification of OGA (L), OGT (M) and OGT/OGA ratio (N) proteins levels in prefrontal cortex from the Western blot in panel B. (O) Western blot of O-GlcNAc levels in prefrontal cortex and hippocampus of male OGT^WT^ and OGT^C921Y^ mice. Actin antibodies were used as loading control. (P) Western blot of O-GlcNAc levels in striatum and cerebellum of male OGT^WT^ and OGT^C921Y^ mice. Actin antibodies were used as loading control.Quantification of O-GlcNAc levels in prefrontal cortex (Q), hippocampus, striatum (S) and cerebellum (T) from the Western blot in panel O and P.).

**Supplementary Figure S14:**
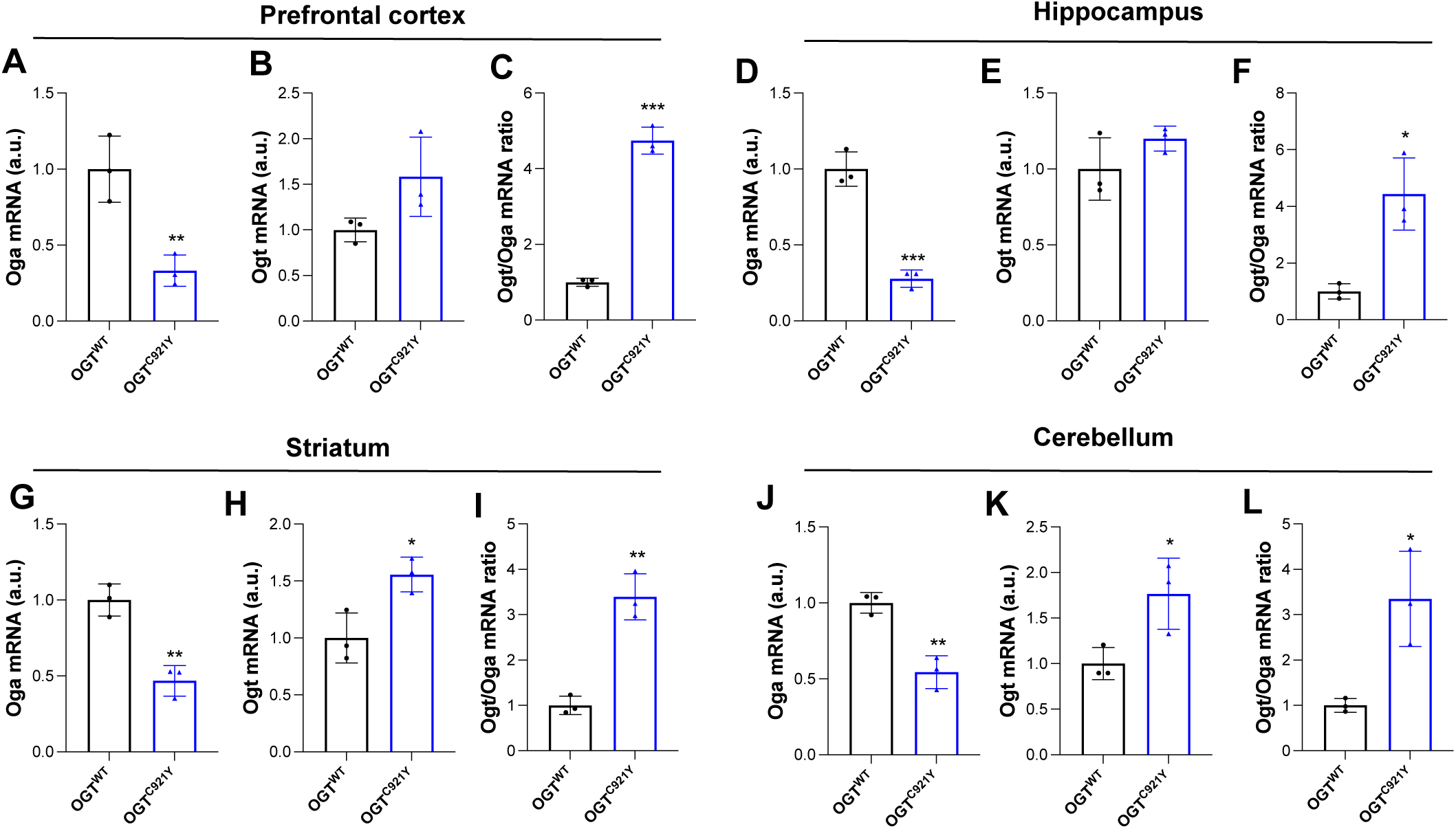
OGT^C921Y^ mice show altered *Ogt*/*Oga* mRNA ratio levels in brain regions. Significance is shown as * *p* < 0.05, ** *p* < 0.01 and *** *p* < 0.001 (n = 3 per group). Student *t* test was used for statistics. Quantification of Oga (A), Ogt (B) and Ogt/Oga ratio (C) mRNA levels in prefrontal cortex Quantification of Oga (D), Ogt (E) and Ogt/Oga ratio (F) mRNA levels in hippocampus. Quantification of Oga (G), Ogt (H) and Ogt/Oga ratio (I) mRNA levels in striatum. Quantification of Oga (J), Ogt (K) and Ogt/Oga ratio (L) mRNA levels in cerebellum.

**Supplementary Figure S15:**
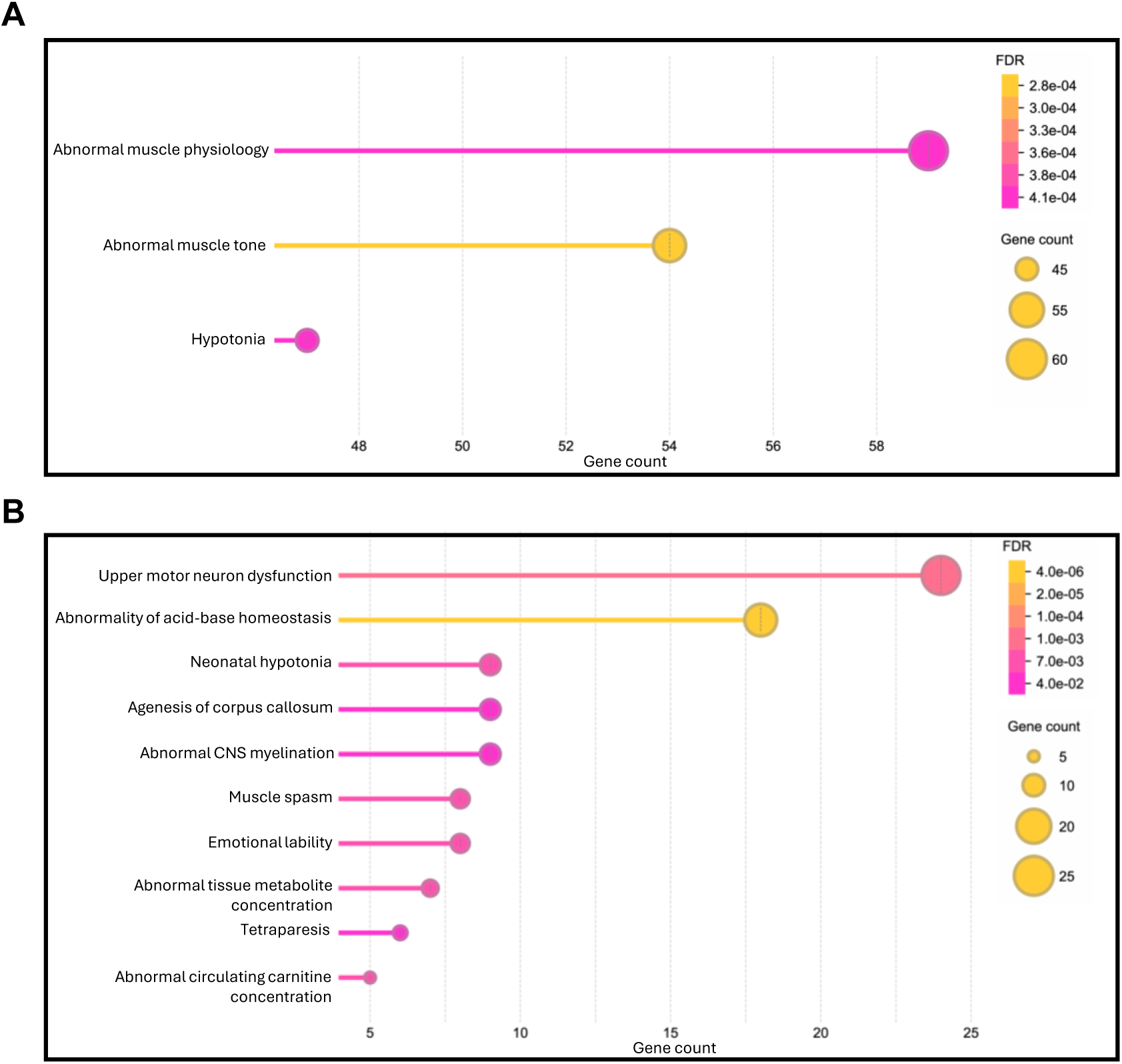
**Perturbed proteomic signatures in the prefrontal cortex corresponded to distinct Monarch enrichment profiles in the OGT^C921Y^ mice** (A) Up-regulated human phenotypes enrichment in the Monarch database. (B) Down-regulated human phenotypes enrichment in the Monarch database.

